# Human NK cell deficiency as a result of biallelic mutations in *MCM10*

**DOI:** 10.1101/825554

**Authors:** Emily M. Mace, Silke Paust, Matilde I. Conte, Ryan M. Baxley, Megan Schmit, Nicole C. Guilz, Malini Mukherjee, Ashley E. Pezzi, Jolanta Chmielowiec, Swetha Tatineni, Ivan K. Chinn, Zeynep Coban Akdemir, Shalini N. Jhangiani, Donna M. Muzny, Asbjørg Stray-Pedersen, Rachel E. Bradley, Mo Moody, Philip P. Connor, Adrian G. Heaps, Colin Steward, Pinaki P. Banerjee, Richard A. Gibbs, Malgorziata Borowiak, James R. Lupski, Stephen Jolles, Anja K. Bielinsky, Jordan S. Orange

## Abstract

Human natural killer cell deficiency (NKD) arises from inborn errors of immunity that lead to impaired NK cell development, function or both. Through the understanding of the biological perturbations in individuals with NKD, requirements for the generation of terminally mature functional innate effector cells can be elucidated. Here we report a novel cause of NKD resulting from compound heterozygous mutations in MCM10 that impaired NK cell maturation in a child with fatal susceptibility to CMV. MCM10 has not been previously associated with monogenic disease and plays a critical role in the activation and function of the eukaryotic DNA replisome. By modeling MCM10 deficiency in human NK cell lines and primary NK cell precursors, we demonstrate that MCM10 is required for NK cell terminal maturation and acquisition of immunological system function.

## Introduction

Human natural killer (NK) cells play a critical role in the control of viral infection and malignancy through contact-mediated killing of susceptible target cells and cytokine secretion. While rare, monogenic cases of human NK cell deficiency (NKD) lead to severe and life-threatening illness in which the abnormality of NK cells is the major clinically relevant immunological defect (reviewed in (1–4)). To date, five causes of classical or developmental NKD, in which loss of NK cells or aberrant NK cell development leads to absence of impaired maturation of peripheral blood NK cells, have been described (3). These include GATA binding protein 2 (GATA2), interferon regulatory factor 8 (IRF8), minichromosome maintenance 4 (MCM4) and go-ichi-ni-san complex subunit 1 (GINS1) deficiencies (5–9). In addition, although only reported in one child, regulator of telomere elongation helicase (RTEL1) mutation has been described to be able to cause NKD (10, 11). Each of these cases have revealed previously unidentified requirements for human NK cell differentiation and homeostasis and are distinct from a broader category of inborn defects of immunity that can include NK cell abnormalities as a minor immunological defect. The biology uncovered through discovery of NKD includes a requirement for the CDC45-MCM2-7-GINS (CMG) complex in NK cell terminal maturation, illustrated by patients with MCM4 and GINS1 deficiencies. Here we describe mutations in another factor required for CMG function, minichromosome maintenance 10 (MCM10), that caused NKD in a child with severe and ultimately fatal CMV infection.

Human NK cells develop from the common lymphoid progenitor and undergo increasingly restricted stages of lineage commitment leading to terminal maturation. NK cells comprise 5-20% of the lymphocyte population in peripheral blood and can be further subdivided into two subsets, CD56^bright^ and CD56^dim^, each with unique phenotypic and functional properties. CD56^bright^ NK cells are potent producers of cytokines, particularly interferon-*γ* (12, 13). The majority of NK cells in peripheral blood are CD56^dim^, the subset specialized to mediate target cell lysis. As such, they express perforin and granzymes to lyse susceptible targets and the Fc receptor CD16 to rapidly recognize opsonized cells.

Multiple lines of evidence demonstrate that CD56^bright^ NK cells are direct precursors of CD56^dim^ NK cells, including longer telomeres in CD56^bright^ NK cells and the early appearance of CD56^bright^ NK cells following hematopoietic stem cell transplant (14). Similarly, in models of in vitro differentiation or humanized mouse reconstitution, the CD56^bright^ subset is detected prior to the CD56^dim^ subset, suggesting these are less mature precursors (15—17). Recent single cell analyses of human NK cells include pseudotime projections of gene and protein expression in peripheral blood NK cell subsets that define a spectrum of development ranging from CD56^bright^ to CD56^dim^ (18). However, despite these data from a variety of model systems, the human NK cell developmental program is poorly understood and identifying requirements for NK cell development is an ongoing challenge within the field.

Of the previously described classical NKD, most include an aspect of aberrant NK cell subset generation in addition to reduced NK cell frequencies within the lymphocyte population. A highly conserved feature of the NK cell phenotype in patients with GATA2 deficiency is the near or absolute absence of the CD56^bright^ subset (8, 19–21). CD56^dim^ NK cells may be present in these patients, while their intrinsic function of target cell lysis is impaired and likely contributes to severe herpesviral illness (8). In contrast, IRF8, MCM4 and GINS1 deficiency lead to reduction specifically of the CD56^dim^ subset and concomitant over-representation of the CD56^bright^ subset (5–7, 9). In the case of biallelic IRF8 deficiency, the CD56^dim^ NK cells that are present have impaired lytic function, and therefore it is likely that this impairment combined with reduced CD56^dim^ NK cell frequency leads to unusual susceptibility to viral infections (7). Therefore, the deregulation of NK cell subset generation is frequently accompanied by impaired NK cell function and viral susceptibility.

First described in a cohort of endogamous Irish travelers with familial NKD, a hypomorphic *MCM4* mutation causes susceptibility to viral infections, short stature and adrenal insufficiency (5, 6, 22, 23). Further investigation into the NK cell phenotype in these patients revealed low NK cell numbers with significant over-representation of the CD56^bright^ subset (5, 6). Subsequent to the report of MCM4-deficient patients was the description of individuals with GINS1 deficiency, which leads to a strikingly similar clinical and NK cell phenotype to MCM4 deficiency. In 5 patients from 4 kindreds, compound heterozygous mutations in *GINS1* lead to intrauterine growth retardation, neutropenia and NKD (9, 22, 24). Specifically, low NK cell number in these patients is accompanied by a relative over-representation of the CD56^bright^ subset that is suggestive of the NK cell phenotype in individuals with hypomorphic *MCM4* mutations.

MCM4 and GINS1 are subunits of the CDC45-MCM2-7-GINS (CMG) replicative helicase complex that binds to origins of replication and is required for DNA synthesis. The MCM2-7 complex binds to chromatin in a cell cycle specific manner and is highly expressed in proliferating cells (25). Hypomorphic mutations of *Mcm4* in S. *cerevisiae*, or in the *Mcm4*^Chaso3^ mouse model, lead to genomic instability and increased tumor formation in the mouse (26). While *Mcm4* knockout mice are not viable, mutation of F345I disrupts interactions between MCM family members and leads to cell cycle arrest (26). The reported NKD-causing variants in *MCM4* and *GINS1* are hypomorphic alleles but their effect has not been defined using NK cell experimental systems. Fibroblasts from these individuals have increased genomic in stability and impaired cell cycle progression with increased induction of DNA damage repair pathways (5, 6, 9). The profound NK cell phenotype in these patients, and accompanying viral susceptibility, suggests that the CMG complex is an important regulator of human NK cell terminal maturation, however the mechanism by which this effect is mediated is not well understood.

Here we describe a single patient with unusual susceptibility to CMV infection having NKD. Genetic analyses identified a compound heterozygous mutation in *MCM10*. MCM10 is a highly conserved and non-redundant component of the eukaryotic replisome that binds directly to the MCM2-7 complex, CDC45, and both double-and single-stranded DNA (27). MCM10 is thought to play a critical role in CMG complex assembly and activation and replication fork processivity (28–30). Here, we sought to define the molecular nature of the *MCM10* mutations discovered in a patient with NKD. We also sought to determine the role of MCM10 in human NK cell maturation and function using models of MCM10 knockdown in an NK cell line and primary NK cell precursors. Moreover, we recapitulated NK cell development in vitro and in vivo using patient-derived induced pluripotent stem cells (iPSCs). These studies demonstrate the importance of MCM10 function in human NK cell maturation, accentuate the importance of the CMG to NK cell development, and define MCM10 deficiency as a novel cause of classical NKD.

## Results

### Clinical history and variant allele confirmation

The male proband was born to healthy, non-consanguineous parents yet presented at 16 months of age with fever, organomegaly, diarrhea and CMV infection (2 × 10^6^ copies/ml). T and B cell numbers were slightly decreased with reduction in effector and memory T cells. Profoundly reduced NK cell numbers were noted and further analysis determined that 50% of these were in the CD56^bright^ subset (Fig. 1A, Table 1). T cell activation in response to phytohemagglutinin was reduced relative to control but responses to phorbyl myristate acetate and CD3 activation were normal. While perforin expression was within normal range, elevated levels of ferritin and triglycerides and decreased fibrinogen concentration prompted consideration of hematophagocytic lymphohistiocytosis (HLH). Expression of SLAM-associated protein (SAP), X-linked inhibitor of apoptosis (XIAP), MHC I and MHC II were normal and no mutations in CD3ζ were detected. The patient underwent bone marrow transplantation for suspected PID but succumbed to overwhelming preexisting CMV at 24 months.

**Fig. 1.**
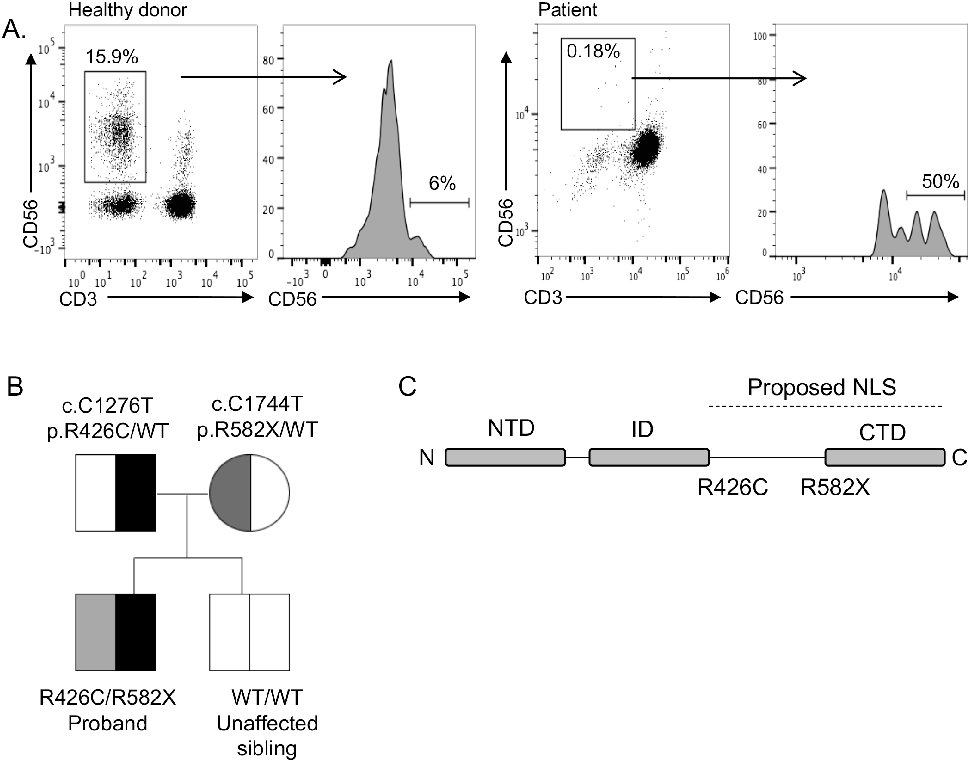
Decreased frequency of peripheral blood NK cells with overrepresentation of the CD56^bright^ subset in an individual with compound heterozygous mutations in MCM10. Severe CMV infection in the male proband born to healthy parents led to evaluation of peripheral blood NK cells and whole exome sequencing of the proband and his immediate family. A) Flow cytometric analysis of peripheral blood NK cells from a representative healthy donor (left) and the proband (right). NK cells are defined as CD56^+^CD3^−^. The relative frequency of CD56^bright^ NK cells within the NK cell subset is defined by density of CD56 staining (histograms). B) Whole exome sequencing identified compound heterozygous mutations that were rare and predicted to be damaging with familial segregation. C) Location of identified variants relative to MCM10 domains. Dashed line indicates previously defined NLS (32). NLS, nuclear localization signal; NTD, N terminal domain; ID, internal domain; CTD, C terminal domain.

**Table 1.** Values from clinical laboratory tests for patient (reference ranges shown in parentheses)

|  |  |
| --- | --- |
| CD3+ | 1250 (1400-8000) |
| CD4+ | 770 (900-5500) |
| CD8+ | 280 (400-2300) |
| CD19+ | 210 (600-3100) |
| CD56+CD3- | 1 (100-1400) |
| CD3+CD8+CD45RA+ | 255 (249-1092) |
| CD3+CD4+CD45RA+ | 855 (455-2454) |
| IgG | 2.22 g/L (3-13.2) |
| IgA | 0.78 g/L (0.3-1.2) |
| IgM | 0.45 g/L (0.48-2.1) |
| Ferritin | 33150 ug/L (15-100 ug/L) |
| Triglycerides | 1.7 mmol/L (0.4-1.6 mmol/L) |
| Fibrinogen | 0.5 g/L (2-4 g/L) |

Exome sequencing (ES) was performed on DNA derived from tissue from the affected individual, his parents, and his healthy sibling. Rare variant, family based genomic analysis of ES results was performed as described previously, filtering on low allelic frequency (31). These analyses identified compound heterozygous mutations in *MCM10* in the affected individual; the variant alleles segregated in accordance with Mendelian expectations for a recessive disease trait and were confirmed by Sanger sequencing (Supp. Fig. 1A).

A missense variant allele in exon 10 was inherited from the father ([NM_018518.4] c.1276C>T, p.R426C) and a nonsense variant introducing a premature stop codon at the end of exon 13 ([NM_018518] c.1744C>T, p.R582X) of this 19-exon gene was inherited from the mother (Fig 1B). This stop-gain variant is located N-terminal to the previously reported C terminal domain required for nuclear localization in multicellular eukaryotes (32) (Fig 1B, C) and is predicted to be subject to degradation by nonsense-mediated decay (33). The missense variant was predicted to be disease-causing by MutationTaster (score 0.99) (34) and likely damaging by PolyPhen2 (score 1.0) (35). The Combined Annotation Dependent Depletion [CADD, http://cadd.gs.washington.edu/] score was 24.3 (36) and the arginine in position 426 is conserved across all species. The nonsense variant was not found in ExAC (37) nor in GnomAD (38), while the missense variant was found at extremely low frequency (4.12 × 10^−5^ in a heterozygous state with no homozygotes in ExAC, 2.5 × 10^−5^ in GnomAD). The rarity and likely damaging effect of these two variants suggesting pathogenic alleles, combined with the clinical and NK cell phenotype of the patient, argued that MCM10 deficiency was responsible for his disease.

### Mutations in *MCM10* affect protein localization and function

To determine the effect of the patient mutations on MCM10 protein expression and function, we performed immunoblotting of lysates from primary dermal fibroblasts obtained from the patient and a healthy, unrelated donor. Western blotting of patient fibroblasts showed the presence of protein at the predicted molecular weight for full-length MCM10, suggesting that the mRNA carrying the premature stop codon is degraded by nonsense-mediated decay and does not permit production of the truncated protein (Fig 2A, Supp. Fig. 2). While the nuclear localization signal (NLS) for human MCM10 has not been specifically defined, it has been localized to the metazoan specific C-terminal region of the protein (32). To understand the effect of C-terminal MCM10 truncations, in the case that they were expressed by the patient (even transiently) stable 293T cells were generated expressing N-terminally tagged full length and truncation mutants. Western blot analyses demonstrated stable accumulation of all tagged versions of MCM10 (Fig. 2B) Furthermore, immunofluorescence experiments showed that tagged MCM10 localizes to the nucleus when truncated C-terminal to the NLS, but remains cytoplasmic when truncated N-terminal to the NLS (Fig. 2C). Given its position relative to the NLS, we hypothesized that if the truncated protein were expressed the MCM10-R582X mutation would prevent its nuclear localization. To specifically test this hypothesis we generated MCM10-turboGFP and turboGFP-MCM10-R582X constructs and transiently overexpressed these in 293T cells. Using confocal microscopy, we found that expression of the R582X nonsense mutation, but not the R426C variant, significantly impaired nuclear translocation of MCM10 (Fig. 2D). Therefore, while endogenous protein in patient cells was not detected due to nonsense-mediated decay, any production of R582X variant demonstrates ineffective nuclear localization due to truncation of the NLS.

**Fig. 2.**
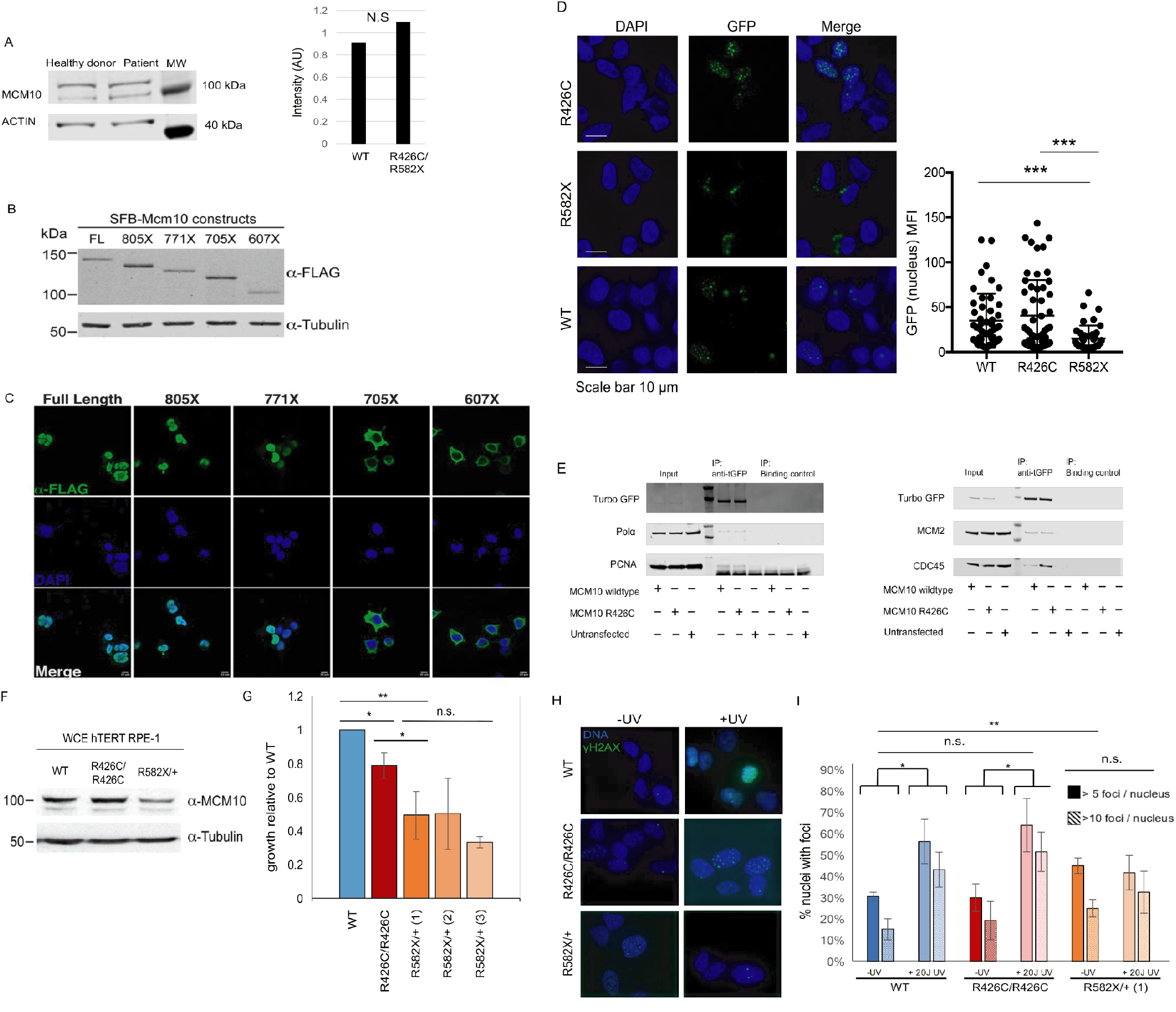
Independent damaging effects of R426C and R582X mutations. A) Primary fibroblasts from the proband and a healthy donor were lysed and probed for MCM10 protein (left). Intensity of MCM10 immunoblot was normalized to loading control (actin, right). B) Western blot analyses of stable SFB-MCM10 expression in 293T whole cell extracts. Full length SFB-MCM10 and truncation mutants were detected with anti-FLAG antibody (top) and with anti-α-tubulin antibody used as a loading control (bottom). C) Confocal imaging of SFB-MCM10 localization in stable 293T cell lines. Full length SFB-MCM10 and truncation mutants were detected with anti-FLAG antibody (green) with DNA indicated by DAPI staining (blue). Merged images show the amount of nuclear SFB-MCM10. D) Wild-type GFP-MCM10 or GFP-R582X MCM10 were transiently expressed in 293T cells. Immunofluorescence by confocal microscopy showing impaired nuclear localization of R582X MCM10 with quantification measuring GFP intensity in the nucleus (right). E) Wild-type MCM10-GFP or R426C MCM10-GFP constructs were transiently expressed in 293T cells. GFP was immunoprecipitated following cell lysis and DNA digestion and blots were probed for POLA, MCM2, PCNA, CDC45. F) Whole cell extract (WCE) of parental (WT), R426C homozygous patient mutation (R426C/R426C) and R582X heterozygous patient mutation (R582X/+) hTERT-RPE-1 cell lines were probed for MCM10 levels. α-tubulin was used as a loading control. G) Parental (WT), R426C homozygous patient mutation (R426C/R426C) and 3 R582X heterozygous mutation clones (R582X/+ 1, 2 and 3) were plated in a 6 well plate, collected and counted after 72 hrs. Doubling time of mutant cells was compared to parental. Error bars represent standard deviation between three independent experiments. Significance for experiment was determined by T test. p<0.05 and p<0.01 are indicated by * and ** respectively. H) Phosphorylated histone variant histone H2AX was used as a marker for DNA damage. I) Foci were counted for a minimum of 30 nuclei in cells treated with 20J UV or untreated. Error bars represent standard deviation between three independent experiments. Significance for experiment was determined by T test. p<0.05 and p<0.01 are indicated by * and ** respectively.

The *MCM10* c.1276C>T, p.R426C allele was predicted to be potentially damaging but not affect protein localization or stability by PolyPhen2 prediction software (35). Expression and localization of WT and MCM10 R426C were comparable when examined using a bicistronic vector expressing full-length WT MCM10 with a GFP reporter in concert with MCM10 R426C with an mApple reporter (Supp. Fig. 3). The use of the bicistronic system also demonstrates stability of the R426C variant relative to WT MCM10, which we could not otherwise assume.

In budding yeast, Mcm10 plays a role in loading and stabilizing Pol-*α* at origins of replication (39–41). Furthermore, direct interaction of di-ubiquitinated MCM10 with PCNA is required for DNA elongation (42). Human MCM10 interacts directly with the CMG helicase through MCM2 and CDC45, supporting chromatin association of MCM10 and efficient firing and elongation during replication (28, 30). To further probe the effect of the R426C mutation on repli-some formation, we immunoprecipitated WT MCM10 or MCM10 R426C using turboGFP and probed for POLA and MCM2, CDC45 and PCNA. The R426C variant did not affect the quantity of POLA, MCM2 and PCNA co-immunoprecipitated with MCM10, and we found a reproducible but insignificant increase of CDC45 binding to the R426C variant compared to WT MCM10 (Fig. 2E). These data suggest that the R426C mutation does not impair formation of the replisome.

To further understand the effects of the patient mutations separately we generated HTERT-RPE-1 cell lines homozygous for the R426C mutation or heterozygous for the R582X mutation. Homozygous lines for the R582X mutation could not be generated as this is presumably a lethal null allele given that MCM10 function is essential. Doubling time was assessed and compared to parental HTERT-RPE-1 cells.The R582X heterozygous mutation had a 50% reduction in MCM10 protein expression as measured by western blot which was accompanied by 50% reduction in growth in 3 unique clones (Fig. 2F). The R426C homozygous cell line retained approx imately 80% growth compared to WT, which is consistent with this mutation having a critical role in the patient’s phenotype, particularly when paired as a compound heterozygous allele with the more severe nonsense mutation (Fig. 2G). This suggests that in human cell lines, expression of the heterozygous R582X mutation leads to MCM10 haploinsufficiency. Furthermore, when assessed for basal levels of DNA damage, the R582X heterozygous cell line had increased levels of *γ*H2AX foci without UV treatment (Fig. 2H). The parental and R426C homozygous cell lines had similar basal DNA damage which significantly increased upon treatment with 20J UV light. However, despite the R582X heterozygous cell line having elevated basal DNA damage, this did not significantly increase following UV treatment (Fig. 2I). Thus, R582X haploinsufficiency accentuates the damaging nature of the R426C allele while not causing a complete absence of MCM10 function, which would be incompatible with cell survival.

To specifically determine the effect of the patient *MCM10* alleles in patient cells, SV40 large T antigen transformation of primary dermal fibroblasts from the proband and an unrelated healthy donor was performed. Western blotting of fractionated cell lysates confirmed the presence of MCM10 in the nuclear fraction of patient-derived cells with slightly increased intensity relative to healthy controls (Fig. 3A). In human cells, MCM10 is released from chromatin at the end of S phase of the cell cycle (40). Given the increased intensity of MCM10 staining in the nucleus of patient-derived cells, we sought to further biochemically quantify the affinity of MCM10 binding to chromatin. We performed tight chromatin fractionation with differential salt extraction. At increased NaCl concentration (0.3M), we found MCM10 more tightly associated with chromatin in patient-derived cells when compared to healthy donor control (Fig. 3B). Given that human MCM10 is normally released from DNA at the end of S phase (40), increased chromatin association in patient cells is suggestive of aberrant cell cycle dynamics and therefore function of the complex containing variant MCM10. Taken together, we demonstrated the impact of each mutation in isolation, including undetectable expression of the R582X variant at endogenous levels and impaired release from chromatin of the R426C variant which is accompanied by impaired growth in a non-transformed human cell line.

**Fig. 3.**
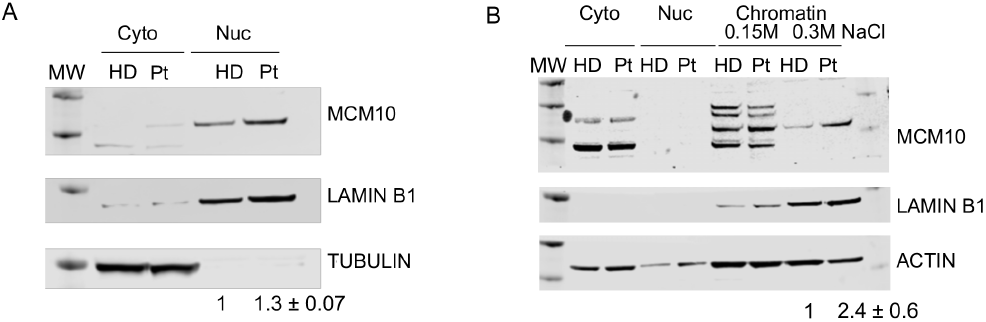
Effect of compound heterozygous mutations. Primary fibroblasts from the patient were generated by SV40 large T antigen transformation. A) Cells were fractionated as described in Methods and nuclear and cytoplasmic fractions were probed for MCM10, LaminB1 and α-tubulin. B) Chromatin fractionation was performed with increasing stringency of salt concentrations (0.15M, 0.3M) from patient fibroblasts immortalized by SV40 large T antigen transduction. Lysates were probed for MCM10, or Lamin B1 and actin as a loading control. Intensity of bands from 0.3M condition are quantified relative to loading control. Values shown are mean ± SD from 3 independent experiments. MW, molecular weight marker; HD, healthy donor; Pt, patient.

### Patient cells have increased nuclear area and frequency of *γ*H2AX foci

Genomic integrity and repair of DNA damage requires CMG helicase function. The previously reported hypomorphic mutations in *MCM4* and *GINS1* lead to abnormal nuclear morphology and increased genomic instability defined by metaphase breaks following aphidicolin treatment (5,9). *GINS1* mutations also lead to increased presence of DNA damage markers, including *γ*H2AX and 53BP1 foci, in primary fibroblasts from patients (9). Using confocal microscopy we visualized the presence of *γ*H2AX foci in immortalized fibroblasts from the patient and a healthy donor (Fig. 4A). We found a significantly higher frequency of *γ*H2AX foci in patient cells, including increased intensity and area of 7H2AX signal within the nucleus of patient cells (Fig. 4A, B). Counter-staining nuclei with DAPI also revealed increased nuclear area in patient cells (Fig. 4A, C), suggesting dysregulation of cell cycle and increased replication stress in patient cells similar to that described for GINS1 deficient primary fibroblasts (9). Transient overexpression of the single mutants in 293T cells led to increased nuclear staining as a result of expression of the R426C variant but not the R582X variant, further supporting the hypothesis that reduced CMG helicase function as a result of the R426C variant leads to replication stress and that this may be manifested as increased nuclear size (Fig. 4D). Finally, we performed cell cycle analyses on immortalized patient fibroblasts with BrdU and 7-AAD. Patient cells had an increased number of cells found in S phase with an accompanying decrease of cells found in G2/M (Fig. 4E). Taken together, these data suggest cell cycle defects and replication stress in patient-derived cells expressing both MCM10 variants.

**Fig. 4.**
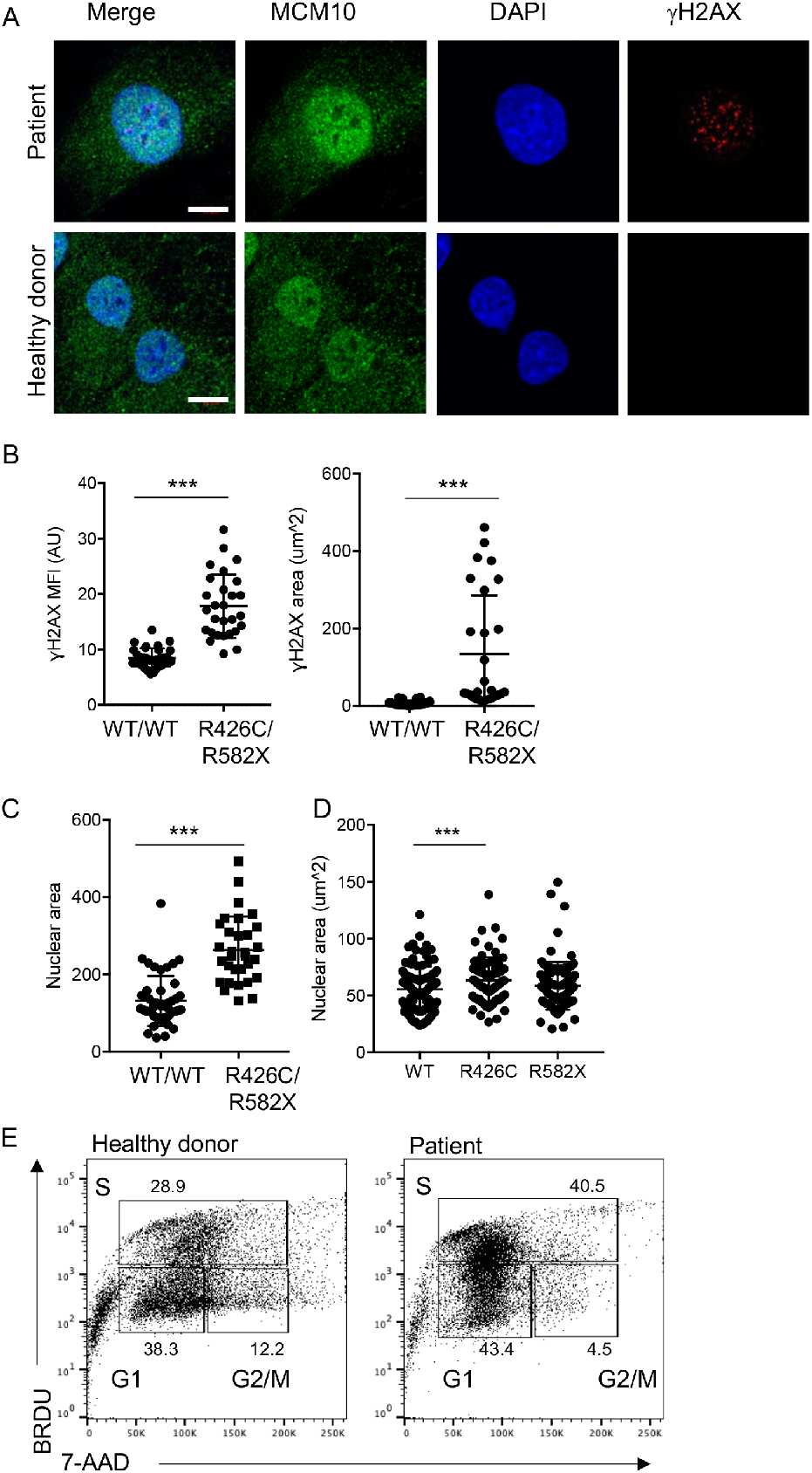
R426C/R582X mutations lead to increased nuclear area and *γ*H2AX staining in immortalized fibroblasts. A) Immortalized fibroblasts were fixed, permeabilized and incubated with primary anti-MCM10 antibody followed by goat antimouse Alexa Fluor 488 secondary antibody and directly conjugated anti-7 H2AX Alexa Fluor 647. Slides were mounted with Prolong Gold antifade media with DAPI and imaged by confocal microscopy. B) MFI of *γ*H2AX staining (left) and area of positive *γ*H2AX signal (right) were measured in >28 cells per condition. Results are representative of 3 independent experiments. C) Nuclear area was measured by positive DAPI staining in 32 (patient) and 46 (healthy donor) cells per condition. Representative of data from 3 independent experiments. ***p<0.0001, Student’s T-test. D) R426C or R582X variants were transiently overexpressed in 293T cells and prepared for microscopy as described above. Nuclear area determined by DAPI staining was measured. Representative of data from 3 independent experiments. *p<0.05, Ordinary one-way ANOVA with multiple comparisons. n=69 (R426C), 95 (R582X, WT). E) Healthy donor-derived (left) or patient-derived (right) immortalized fibroblast cells were labeled with BRDU and 7-AAD and cell cycle was analyzed by FACS. Shown is one representative experiment of 3 independent replicates.

### Impaired cell cycle progression in MCM10 knockdown NK cells

The effect of the R426C and R582X variants in patient cells was suggestive of impairment of CMG helicase function as a result of MCM10 mutation. To model MCM10 deficiency in an NK cell line using a complementary approach to patient cell analyses, we performed CRISPR-Cas9 gene editing of MCM10 in the NK92 human NK cell line. Intronic guides were designed to reduce but not completely abolish protein expression, as cells in which MCM10 was deleted were not viable (not shown). Single cell clones were isolated, expanded and mRNA and protein expression were measured. One clone with approximately 10% protein expression of the parental line was primarily chosen for further study (clone 1), 3 clones were verified with reduced protein expression accompanied by functional defects (Fig 5A, B, not shown). To determine the functional effect of MCM10 knockdown we performed cell cycle analyses in NK92 and NK92 MCM10-knockdown (MCM10-KD) cell lines. These revealed a significant increase in the frequency of cells in early S phase with a decrease of cells in G2/M when compared to the control cell line (Fig. 5C, D). In addition, MCM10-KD cells had significantly increased doubling time (39.7 hrs vs. 28.7 hrs) (Fig. 5E). Thus, the cell cycle profile of MCM10-KD NK92 cell lines was reminiscent of that seen in patient cells and suggested impairment in cell cycle progression, with a specific accumulation of cells detected at early S phase owing to a reduction in MCM10 function.

**Fig. 5.**
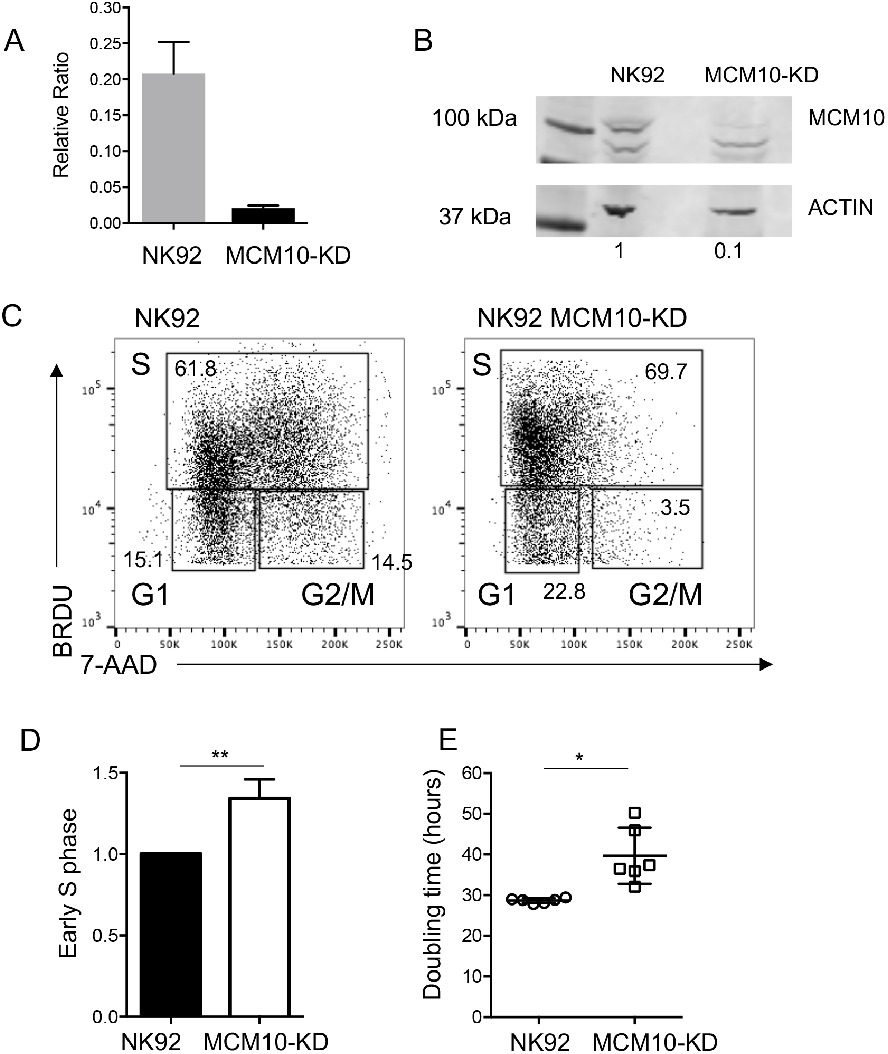
Cell cycle arrest in a cell line model of MCM10 knockdown. MCM10 expression in NK92 cells was reduced by CRISPR-Cas9 gene editing as described in Methods. A) RNA was isolated from wildtype NK92 or MCM10-KD cells and qPCR measurement of MCM10 mRNA was performed. Pooled data from 4 independent experiments done in quadruplicate are shown. B) WT and MCM10-knockdown (KD) NK92 cells were lysed and probed for MCM10 protein and actin as a loading control. Quantification of MCM10 protein relative to loading control is shown below. Data representative of 3 independent repeats. C) WT and MCM10-KD NK92 cells were labeled with BRDU and 7-AAD and cell cycle was analyzed by FACS. D) Quantification of the frequency of cells in early S phase relative to the WT control is shown from 3 independent experiments. E) Cell doubling time was calculated by enumerating cells in culture. Data is pooled from 6 independent experiments.

### MCM10 knockdown leads to replication stress but does not impair cytotoxicity

To further define the role of MCM10 in NK cell genomic integrity, we determined the effect of MCM10 knockdown on indicators of replication stress in the NK92 cell line. NK92 MCM10-KD cells line had greater number and intensity of *γ*H2AX foci at baseline as measured by confocal microscopy (Fig 6A-C).

**Fig. 6.**
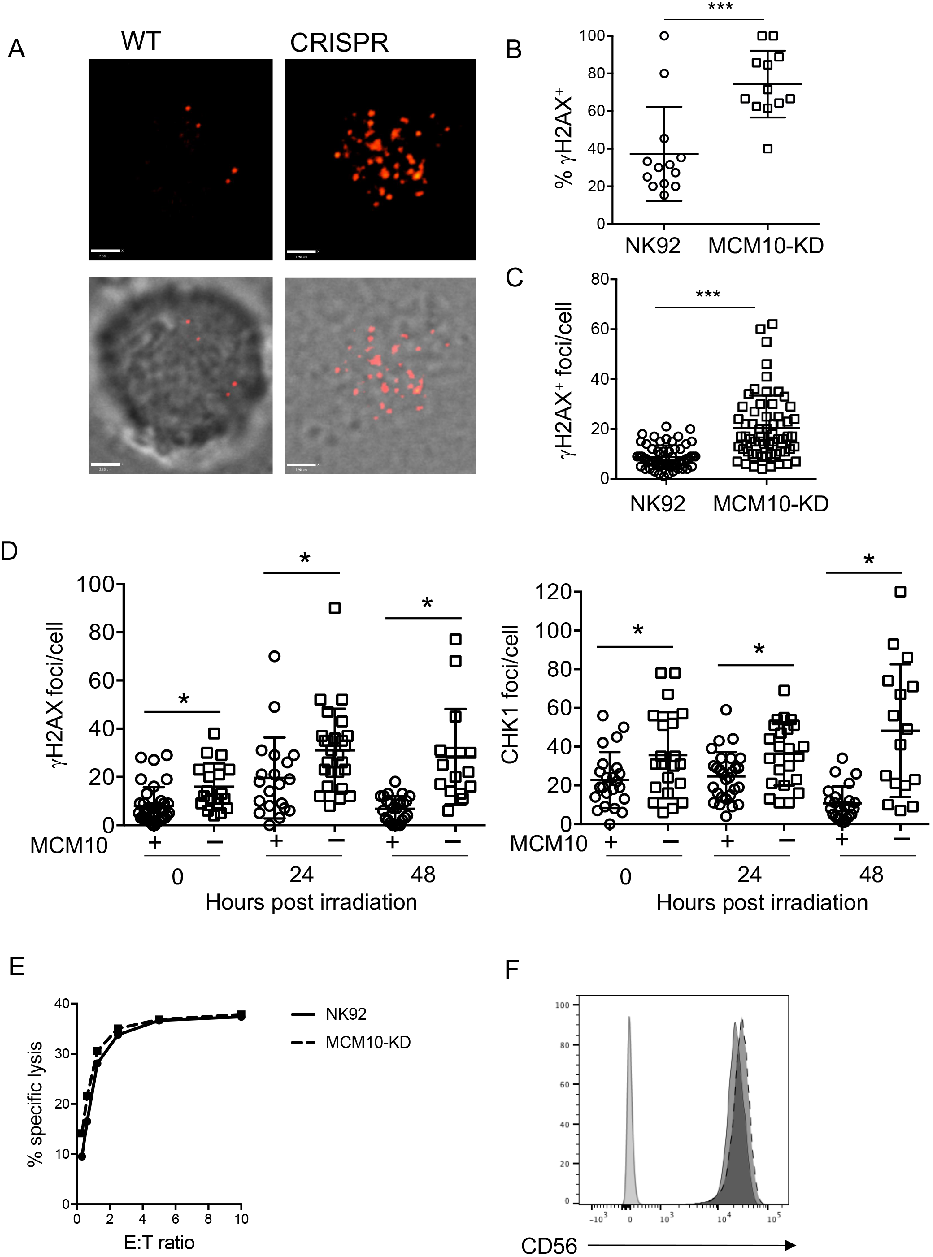
Increased replication stress in an MCM10-KD cell line. WT or MCM10-KD NK92 cells were fixed, permeabilized and incubated with anti-*γ*H2AX Alexa Fluor 647. A) Images were acquired by confocal microscopy. Scale bar 1.9 μm. B) The frequency of cells per field of view that were positive for *γ*H2AX by microscopy was scored by manual counting. ***p=0.0003, n=12. C) The number of *γ*H2AX foci per cell were counted by manual counting of microscopy images. ***p<0.0001, n=60 from 3 independent experiments. D) WT NK92 (+) or NK92 MCM10-KD (-) were irradiated with 2 Gy and then allowed to recover for times indicated prior to fixing and immunostaining for *γ*H2AX (left) or phospho-CHK1 (right). Images were acquired by confocal microscopy and number of foci were enumerated by manual counting. *p<0.05, data pooled from 3 independent repeats. E) Cytotoxic function of wild-type NK92 cell line or MCM10-KD cells against K562 targets was performed by 51Cr release assay. Representative shown from 3 independent experiments. F) CD56 expression on NK92 (solid line) or MCM10-KD (dashed line) NK92 cells was measured by FACS analysis. Data representative of 3 independent repeats. *p=0.03 by Student’s T test. **p=0.08 by Student’s T test.

To further test response to genotoxic stress, cells were irradiated and then rested for 24 or 48 hours before quantitative analysis of *γ*H2AX and phospho-CHK-1, an indicator of ATR pathway activation, by confocal microscopy. This showed a significant increase in the number of *γ*H2AX foci at baseline in MCM10-KD cells, as previously shown (Fig. 6C), as well as in response to irradiation (Fig. 6D). The number of foci increased over time and remained significantly different at 24 and 48 hours post-irradiation when compared to parental cells. Similarly, the levels of phospho-CHK-1 foci per cell was significantly increased in MCM10-KD cells in the absence of irradiation and both 24 and 48 hours postirradiation (Fig. 6D).

The severe CMV infection in this patient suggested impaired NK cell cytotoxic function, however this was not tested due to the limited availability of peripheral blood from the deceased patient. To determine whether NK92 lytic function was directly affected by MCM10 knockdown in the NK92 cell line we performed 4-hour 51Cr release assays and found no difference between WT and MCM10-KD cells when cells were seeded at equal densities overnight prior to the assay to approximate cell cycle synchronicity (Fig. 6E). Furthermore, expression of NK cell receptors, including CD56, were unaffected (Fig. 6F). Thus, the roles of MCM10 in promoting effective replication and cell cycle progression defined in other cells can be extended to NK cells specifically. Further, the impact on primary NK cell function in the patient seemingly arises from aberrant development and not from a direct role for MCM10 in regulating cytotoxic function.

### In vitro differentiation of MCM10-KD NK cells from CD34^+^ precursors

While there was limited biological material available from our patient, flow cytometry of peripheral blood lymphocytes demonstrated profoundly reduced frequency of NK cells, with increased representation of the CD56^bright^ subset. This was analogous to the NK cell phenotype seen in patients with *MCM4* and *GINS1* mutations (6, 9), however the mechanism leading to this phenotype in any of these disorders is unknown. CD56^bright^ NK cells are thought to be the precursors of the CD56^dim^ subset, and previous studies suggest that hyperproliferation of the CD56^bright^ subset is required for their terminal maturation (5, 9, 43). However, whether this defect occurs over time and reflects impaired NK cell homeostasis in the periphery or occurs during differentiation has been untested.

To determine the effect of MCM10 knockdown on NK cell terminal maturation, we performed CRISPR-Cas9 gene editing of CD34^+^ hematopoietic stem cells (HSC) from a healthy donor to generate MCM10-KD precursors, which were used for the generation of mature NK cells by in vitro differentiation (7, 8, 44). CD34^+^ HSC were isolated by FACS from apheresis product and the CRISPR-Cas9 guide found to be most effective for the NK92 cell line or an empty vector containing GFP were nucleofected into purified HSC. These were expanded for 3 days in TPO, IL-3, IL-7 and SCF, re-sorted to select GFP^+^ cells, and were then cultured in vitro in the presence of EL08.1D2 stromal cells to generate mature NK cells. Following 30 days of culture, cells were harvested and NK cell developmental stages were assessed by flow cytometry. NK cells derived from CD34^+^ HSC transfected with empty vector underwent terminal maturation with increased frequency when compared to those transfected with MCM10 CRISPR guide constructs. Specifically, control NK cells up-regulated CD16, whereas MCM10-deficient NK cells showed decreased CD16 signal (Fig. 7A). Developmental subsets were quantified using previously described phenotypic markers that mark progression from Stage 1 (CD34^+^ CD117^−^ CD94^−^ CD16^−^) through Stage 2 (CD34^+^CD117^+^CD94^−^CD16^−^), then Stage 3 (CD34^−^CD117^+^CD94^−^CD16^−^), Stage 4 (CD34^−^CD117^+^/-CD94^+^CD16^−^) and finally Stage 5 (CD34^−^CD117^−^CD94^+^/^−^CD16^+^) (45). There was an increase in Stage 3/early Stage 4 cells in MCM10-KD cells when compared to mock transfected cells. As suggested by the reduced CD16 expression on MCM10-KD cells, the increase in Stage 3 cells was accompanied by a decrease in more mature Stage 4 cells and Stage 5 NK cells when compared to control cells (Fig. 7B). In addition, a subset of cells remained in Stage 2 in MCM10-KD conditions, whereas there were no detectable Stage 2 cells in the control condition. Taken together, these data demonstrate that decreased expression of MCM10 leads to the NK cell phenotype observed in the peripheral blood of a patient with MCM10 mutations. The effect is therefore specific to MCM10 and not a feature of any other genetic influences in the patient as the developmental abnormality can be recreated in healthy donor cells. Importantly, this distinctive phenotype can be recapitulated in vitro and suggests that progression through maturation is impaired at multiple stages of NK cell development and that intact MCM10 function is needed for NK cell maturation.

**Fig. 7.**
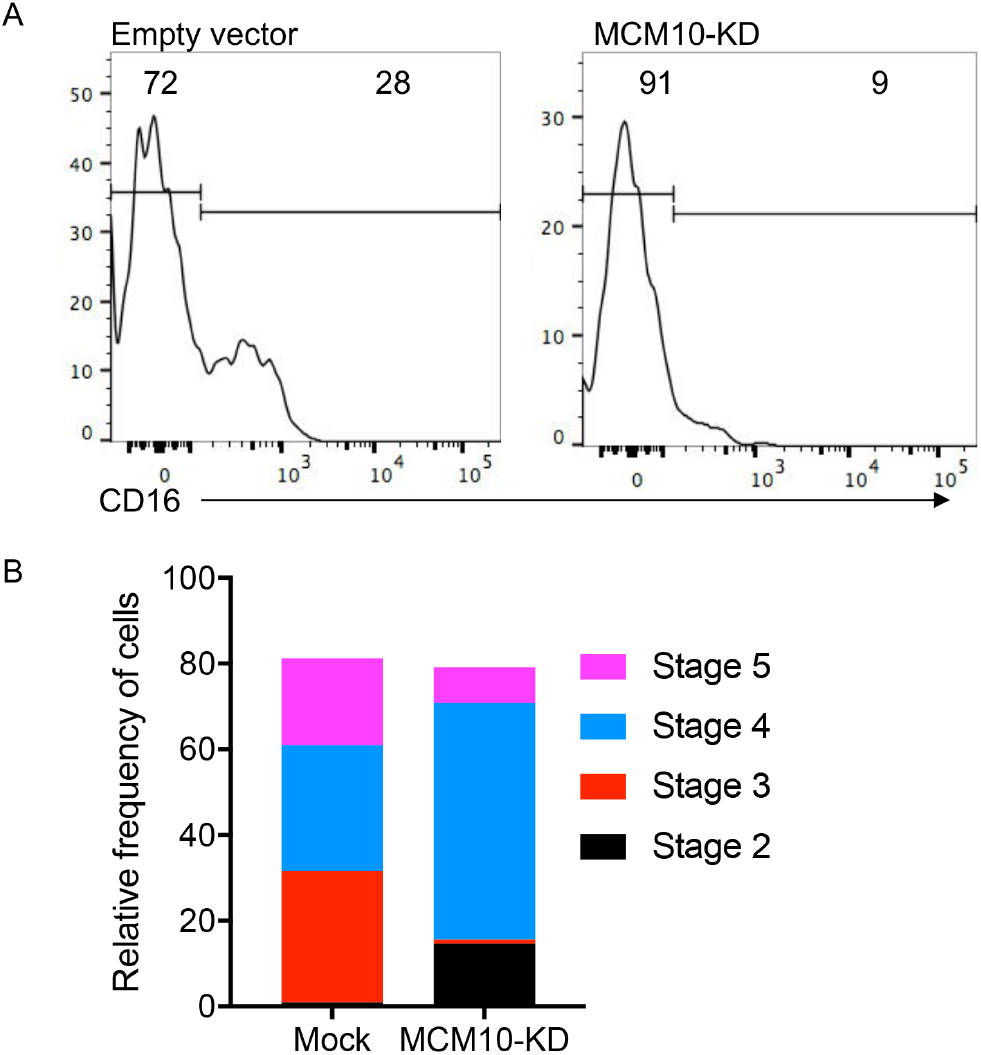
MCM10-KD in primary cells impairs NK cell maturation from CD34^+^ hematopoietic stem cells. CD34^+^ HSC precursors were isolated from apheresed peripheral blood and transfected with MCM10 CRISPR-Cas9-GFP. After 3 days of expansion GFP-positive or-negative cells were sorted and then co-cultured with EL08.1D2 stromal cells in the presence of cytokines, including IL-15, as described in Methods. Cells were harvested at 28 days and NK cell maturation was analyzed by FACS. A) Representative histograms of CD16 expression as a marker of NK cell terminal maturation. B) Relative frequency of cells according to defined stages of NK cell maturation; Stage 1 (CD34^+^ CD117^−^ CD94^−^ CD16^−^), Stage 2 (CD34^+^ CD117^+^ CD94^−^CD16^−^), Stage 3 (CD34^−^CD117^+^ CD94^−^CD16^−^), Stage 4 (CD34^−^CD117^+/−^CD94^+^ CD16^−^) and Stage 5 (CD34^−^CD117^−^CD94^+/−^CD16^+^). Representative of 2 independent experiments.

### Generation of patient-derived NK cells in a humanized mouse model

Finally, we sought to definitively link patient mutations with the NK cell developmental phenotype generated in vitro and observed in the limited ex vivo patient samples. While there were no remaining preserved peripheral blood cells from the deceased patient, we reprogrammed primary patient fibroblasts using a footprint-free modified RNA method to generate iPSC lines. An iPSC line from a healthy donor was generated in parallel. We found both patient and control iPSC lines to have normal karyotypes, express pluripotency markers and undergo differentiation into three-germ layers (not shown).

To test the effect of patient mutations on NK cell development in an in vivo model, we reconstituted humanized mice with CD34^+^ precursors generated from patient or healthy donor iPSCs. For production of CD34^+^ cells, teratomas were generated, harvested and purified. These were injected into NSG mice along with OP9w3a feeder cells and were allowed to develop for 21 days. Tissues were harvested and evaluated by flow cytometry using a human NK cell developmental marker phenotype panel. Notably, this showed significant enrichment of the CD56^bright^ subset within engrafted human NK cells in the blood and spleen from all 4 patient-generated mice (Fig. 8A, B). Further, confocal microscopy showed an increase in the number of *γ*H2AX foci in enriched human NK cells from mice reconstituted with patient-derived iPSCs (Fig. 8C), reminiscent of the increase in foci seen in NK92 MCM10-KD and MCM10 variant-containing cell lines. This was significantly greater when all 4 mice were taken into account (Fig. 8D). Therefore, using complementary mechanisms of modeling patient mutations and MCM10 insufficiency, we demonstrate that MCM10 function is required for the generation of terminally mature NK cell subsets both in vitro and in vivo and that biallelic variants can result in a human NKD.

**Fig. 8.**
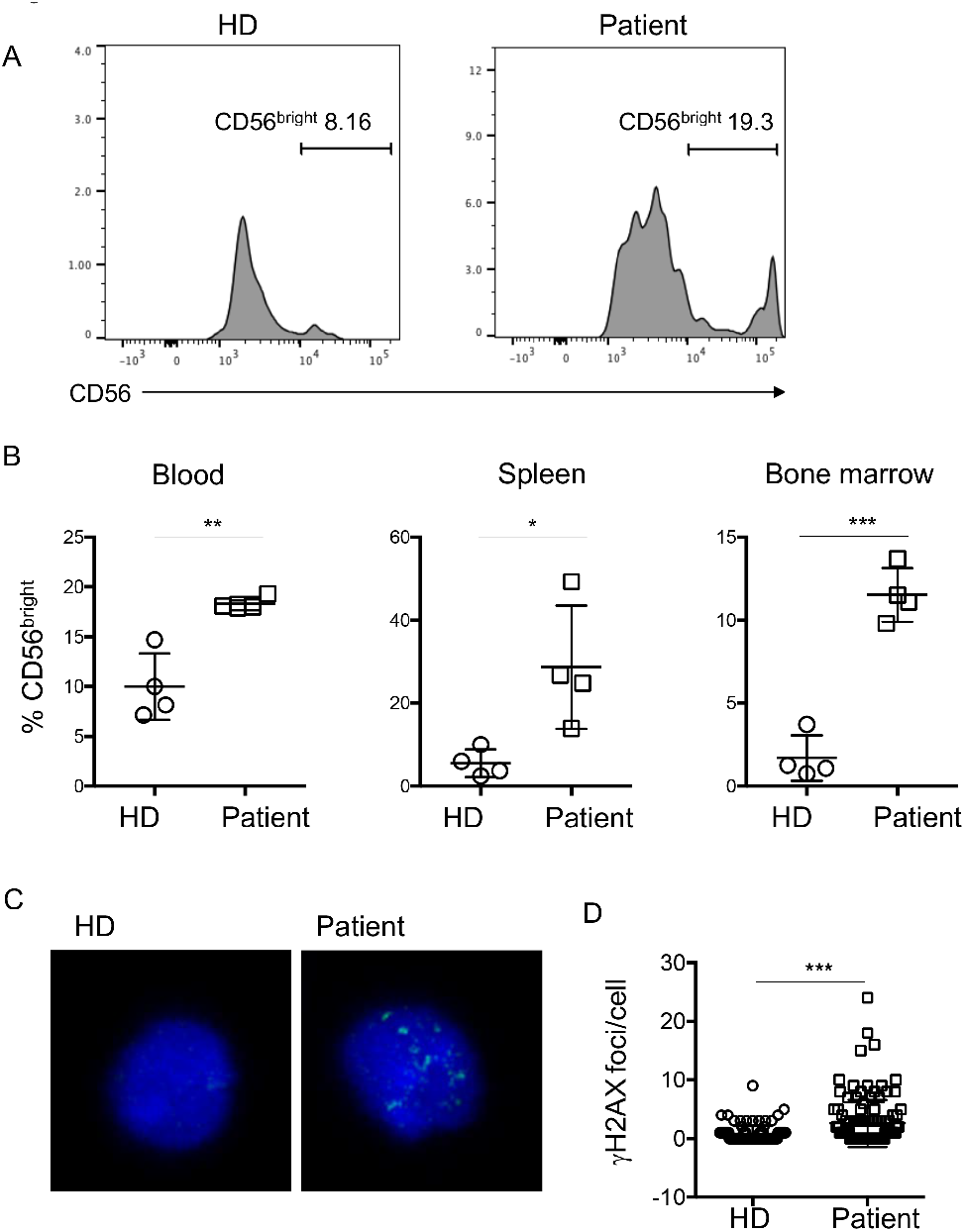
iPSC-derived NK cells from the proband have impaired terminal maturation and increased replication stress. iPSCs reprogrammed from patient primary fibroblasts were differentiated by teratoma formation to CD34^+^ HSC, purified, and transplanted into NSG mice as described in Methods. Organs were harvested 21 days following transplantation and human CD45^+^ CD56^+^CD3^−^ cells were analyzed for density of CD56 expression. n=4 mice per genotype (patient and healthy donor derived iPSCs). A) Representative FACS histograms of NK cells from bone marrow of mice reconstituted with human NK cells from healthy donor (HD) or patient-derived CD34+ cells generated from iPSCs. B) Frequency of CD56^bright^ NK cells from 4 mice per genotype from blood, spleen, and bone marrow as indicated. NK cells were identified as human CD45^+^ CD56^+^CD3^−^ (bone marrow: 93-979 NK cells, blood: 20-405 NK cells, spleen: 60-880 NK cells, all from approximately 10^6^ cells from each organ per mouse) and the frequency of CD56^bright^ NK cells based on CD56 density within the human NK cell population are shown. C) Splenocytes from mice transplanted with healthy donor (HD) or patient-specific iPSC derived CD34+ cells were fixed, permeabilized and incubated with anti-*γ* H2AX antibody. Images were acquired by confocal microscopy. D) Frequency of *γ*H2AX foci per cell were enumerated by manual counting of 88 (HD) and 108 (patient) cells. *p<0.05, **p<0.005, ***p<0.0001 by unpaired Student’s T test.

## Discussion

The study of human NKD provides important insights into the requirements for NK cell differentiation and maturation, particularly when NK cells are the primarily affected lymphocyte subset. To date, 5 monogenic causes of NKD affecting development have been described in literature. Notably two of these, *MCM4* and *GINS1* deficiencies, are a result of mutations that destabilize the CMG complex and lead to reduced numbers of peripheral blood NK cells with relative over-representation of the CD56^bright^ and reduced frequency of CD56^dim^ NK cells (5, 6, 9). While other immune subsets, including neutrophils, ILCs and T cells, may also be affected to varying degrees (9), the profound NK cell-intrinsic defects observed in these rare cases and their association with the patient’s clinical phenotype defines these monogenic discorders as NKD.

Here we report a novel classical NKD as a result of compound heterozygous mutations in *MCM10*. By modeling MCM10 deficiency in a human NK cell line we demonstrate that MCM10 function is required for fidelity of cell cycle progression and decreased MCM10 function leads to increased replication stress. In addition, we recapitulate the NK cell phenotype observed in our proband using CRISPR-Cas9-mediated modeling of MCM10 deficiency followed by NK cell in vitro differentiation and a humanized mouse model of NK cell differentiation from patient iPSCs. These approaches show for the first time that impaired function of a eukaryotic replisome complex member can directly lead to the NK cell phenotype observed in peripheral blood of patients with CMG complex mutations, namely reduced NK cell number and increased representation of the CD56^bright^ subset.

The clinical hallmark of NKD is susceptibility to viral infections, particularly those of the herpesviral family, and the remarkable susceptibility of the proband to CMV prompted investigation into NK cell phenotype and function. This evaluation revealed a profoundly reduced frequency of NK cells in peripheral blood (<1%). In addition, the phenotype of NK cells in peripheral blood was reflective of impaired terminal maturation or survival of terminally mature subsets defined by significant over-representation of the CD56^bright^ subset relative to CD56^dim^ NK cells. Exome sequencing of the patient and his immediate family identified variants in *MCM10* which segregated appropriately and were predicted to be highly damaging. The novelty and predicted damage of these mutations, taken together with the consistency of the NK cell phenotype reported previously for *MCM4* and *GINS1* deficiencies (5, 6, 9), made *MCM10* the strongest gene candidate in this patient.

MCM10 is a critical functional regulator of the CMG complex, and here we demonstrate that both the R426C and R582X mutations in concert are damaging (albeit in different ways) and lead to replication stress and impaired cell cycle progression. Specifically, we show that the R582X variant, which introduces a premature stop codon, presumably leads to nonsense-mediated decay. While the mechanistic impact of the R426C mutation is less clear, our data suggest that it affects cell proliferation, possibly by impairing origin firing and/or replication elongation and resulting inefficiency. This is manifested by increased retention of MCM10 on chromatin and impaired cell growth. Together, the effects of both mutations are increased accumulation of cells in early S phase and replication stress indicated by increased phosphorylation of *γ*H2AX. The slightly increased retention of MCM10 on chromatin under high salt conditions when isolated from patient cells is compatible with altered replication dynamics despite intact interactions with the MCM2-7 complex and Polα. While overexpression of the R582X mutant allele demonstrated its nuclear exclusion, we failed to detect truncated protein in lysates from patient cells by Western blotting. This suggests that MCM10 R582X protein is not expressed at endogenous levels. While quantitative measurement of protein levels did not identify differences in expression of full-length protein between the patient and healthy donors, the cell-cycle dependent changes in expression of MCM10 also make it difficult to draw quantitative conclusions based on this measurement. With relative expression of MCM10 highest in S phase (28), it is likely that cell cycle-dependent protein regulation affects the overall expression of MCM10 in patient cells, which we have shown are more frequently detected in S phase than healthy donor control cells.

The addition of MCM10 deficiency as an NKD strengthens the importance of the CMG complex and DNA replicative machinery in human NK cell functional maturation. With highly conserved NK cell and clinical phenotypes, patients with *MCM4*, *GINS1* and *MCM10* mutations have illustrated the importance of the eukaryotic DNA helicase in NK cell development. Despite the effect of these mutations on cell cycle progression and DNA damage response in extra-immune cell types, the most substantive clinical phenotype in these patients is unusually severe viral susceptibility with accompanying NK cell deficiency. CD56^bright^ NK cells from patients with GINS1 or MCM4 mutations fail to proliferate in response to IL-2, whereas the CD56^dim^ subset undergoes proliferation at rates similar to controls from healthy donors (5). In addition, while less well described, the presence of NK cell impairment in patients with *POLE1* (46) and *POLE2* mutations (47) and a single patient with *RTEL1* mutations (10, 11) suggests that these other regulators of DNA replication and damage repair may similarly function to regulate NK cell homeostasis.

It is remarkable that NK cells are seemingly profoundly sensitive to loss-of-function mutations in these proteins, and the outstanding question remains as to why this is the case. Mouse models demonstrate that complete deletion of MCM genes, including *Mcm4* (26) or *Mcm10* (48) leads to embryonic lethality. Hypomorphic alleles of *Mcm4* (*Mcm4*^chaos3/chaos3^), however, lead to defects in cell proliferation and cancer susceptibility (26), and while milder than MCM4 patient phenotypes, decreased frequencies of NK cells are found in these mice (5). Relatively decreased CMG complex expression in stem cells, which occurs with aging, leads to increased replicative stress and decreased ribosomal function (49). Overexpression or haploinsufficiency cause similar defects (50), demonstrating the importance of tuned levels of protein expression in CMG helicase function.

Tightly controlled CMG complex protein expression is linked to the importance of replicative helicase components to both initiate replication and repair stalled forks if necessary (29). Therefore, it may be that cells with a critical requirement for rapid proliferation, including T cells, have higher production of these proteins and thus are more tolerant of loss of function than NK cells. In support of this, both effector T cells and germinal center B cells have significantly higher expression of MCM10 than mature NK cells in peripheral blood (51). The potential for varying availability of functional CMG protein to differentially affect different lymphocyte subsets makes it conceivable that there are more damaging mutations that lead to combined immune deficiency and have not been detected; similarly, these may not be supportive of viability. In addition, the transient T cell lymphopenia observed in GINS1 deficient patients and the impaired T cell differentiation noted amongst our patient’s PBMC further suggest that, depending on the severity of the mutation, there may be variable effects on other lymphocyte subsets. However, we propose that the particular mutations that cause NKD fall within a window that primarily impacts NK cells rendering the NK cell-mediated defense inadequate; thus leading to susceptibility to viral infections, particularly of the herpesvirus family. As such, the NK cell abnormality would be the main clinically relevant immune deficiency. This is supported by the finding that the clinical severity of disease correlates with the relative levels of GINS complex expression in GINS1-deficient individuals (9). Finally, while the immune compartment in these patients is highly affected, there is an impact of these mutations on cell cycle and DNA damage repair pathways in fibroblasts. While some MCM4 patients report malignancies (5), unfortunately many of these patients have succumbed to viral infections at an unusually young age, perhaps obscuring unusual cancer susceptibility.

In the case of MCM4 mutations and, unlike GINS1 deficient patients, neutrophil numbers were normal in our patient. As such, there are subtle differing effects of CMG mutations on both the clinical manifestations and immune cell phenotypes in patients with MCM4, MCM10 and GINS1 deficiencies. Adrenal insufficiency was reported in MCM4 patients, but has not been detected in GINS1 patients or in the patient reported here with MCM10 deficiency (and our patient did not have intrauterine growth abnormalities). It is unclear whether these differences reflect non-redundant roles for CMG complex proteins or whether they are indicative of differing severity of mutations leading to varying effects on different cell lineages. While innate lymphoid cell (ILC) subsets were not examined in our patient due to limited availability of patient material, GINS1, but not MCM4 patients had loss of ILC subsets in peripheral blood, again reflecting either a more conserved requirement for GINS1 in neutrophil and ILC development in addition to that of NK cells, or reflecting a more severe mutational burden that led to this phenotype.

Regardless of their differences, the conserved impact on NK cells is a notable feature of CMG helicase mutations, especially in light of this being the third independent association of NKD with the CMG complex. Given the impaired proliferation and increased apoptosis of the CD56^bright^ subset it seems that impaired generation of the CD56^dim^ NK cell subset in particular is a feature of the NK cells found in peripheral blood of MCM4 and GINS1 deficient patients (5, 9). Here, we have described an additional patient fitting this paradigm but with biallelic MCM10 mutations associated with reduced NK cell numbers in the periphery and overrepresentation of the CD56^bright^ subset. Notably, we show that this particular NK cell phenotype can be recapitulated using in vitro NK cell differentiation from MCM10 knockdown CD34^+^ precursors or the generation of patient NK cells from iPSCs in a novel humanized mouse model. Therefore, either a survival defect of the CD56^dim^ subset is manifested within the relatively short time frame of these experiments, or there is impaired generation of these cells within these systems. Interestingly, the highest expression of MCM complex components is in stem cells, and CD34^+^ HSCs, as well as other progenitors, have significantly increased expression of MCM10 relative to other cells of hematopoietic and immune origin (51). Our data using in vitro differentiation shows that progression through early stages of NK cell development is also impaired, suggesting that the requirement for proliferation is not restricted to later stages of maturation, but limits access to it.

Finally, there may be additional functions mediated by the CMG helicase that contribute to the NK cell phenotype in these patients. MCM5 binds directly to the STAT1α promoter and mediates direct transcriptional control of IFN*γ* response genes (52). Careful genetic and functional analyses of the effect of these mutations on gene expression throughout NK cell development will be important for definitively elucidating the nature of these deficiencies on NK cell maturation and function. In addition, fully understanding the effect of such mutations on adaptive NK cells may shed additional light on the requirement for proliferation in their generation and the role that the adaptive subset plays in viral control.

In summary, here we define a novel classical NKD as a result of biallelic mutations in *MCM10* leading to impaired generation of terminally mature NK cells and loss of NK cell function. While rare, NKD provide invaluable insight into requirements for human NK cell differentiation and function, and the discovery of *MCM10* as an NKD gene strengthens the importance of intact function of the eukaryotic helicase complex in human NK cell functional maturation, with this representing the third such association. In addition, through our modeling of MCM10 insufficiency in NK cell development, we provide important mechanistic insight into the underlying immunological and biological cause of these defects.

## Methods

### Animal studies

NOD.Cg-Prkdc^scid^ Il2rg^tm1Wjl^/SzJ mice were purchased from Jackson Laboratories (Bar Harbor, Maine) and housed in pathogen-free conditions in the vivarium at Texas Children’s Hospital, Houston, TX. All mice were cared for in the animal facilities of the Center for Comparative Medicine at Baylor College of Medicine (BCM) and Texas Children’s Hospital (TCH), and all protocols were approved by the BCM Institutional Animal Care and Use Committee.

### Human study approval

All studies were performed in accordance with the Declaration of Helsinki with the written and informed consent of all participants under the guidance of the Children’s Hospital of Philadelphia, Baylor College of Medicine, Columbia University and University Hospital Wales.

### Exome sequencing and analysis

Exome sequencing was performed on genomic DNA as part of the primary immunodeficiency cohort study at Baylor Hopkins Center for Mendelian Genomics (BHCMG, http://bhcmg.org/) (31, 53). Exonic portions of genomic DNA were captured with custom VCrome probes (52 Mb; Roche NimbleGen Inc.) and DNA was sequenced using Illumina HiSeq 2500 equipment (Illumina). Sample preparation, capture method, sequencing, bioinformatic processing, filtering and the gene variant databases used, and further manual evaluation and interpretation of the exome data have previously been published in detail (53). Bioinformatic analyses were performed as previously described using filters of an allelic frequency of less than 0.0005 in the Baylor Hopkins Center for Mendelian Genomics (BHCMG, http://bhcmg.org/) (54), ESP5400 data of the National Heart, Lung, and Blood Institute GO Exome Sequencing Project (http://evs.gs.washington.edu/EVS), 1000 Genomes (http://www.internationalgenome.org/) (55) and ExAC (http://exac.broadinstitute.org/) (37) databases. Variants identified were subsequently confirmed by Sanger dideoxy DNA sequencing using primers constructed by Primer3 (56). Results from Sanger sequencing were visualized using 4Peaks software (Nucleobytes).

### Cell isolation, cell lines and transfection

Primary peripheral blood mononuclear cells were isolated by Ficoll-Paque density centrifugation. CD34^+^ cells for in vitro differentiation were isolated from discarded apheresis product from routine red cell exchange at Texas Children’s Hospital. NK92 cells were maintained in Myelocult medium supplemented with penicillin/streptomycin and 100 U/ml IL-2 (Roche). CRISPR-edited cell lines were generated by nucleofecting NK92 cells or CD34^+^ precursors with pCMV-Cas9-GFP all-in-one plasmid (Sigma-Aldrich) with the intronic guide sequence GAT TTA ATA TTC CCC GCT TGG G. Single cell clones were isolated by FACS sorting for GFP positive clones and expanded in culture. MCM10 knockdown was measured by Western blot and normalized to a loading control as described below. Primary patient or healthy donor fibroblasts were immortalized by lentiviral transduction with SV40 large T antigen transformation, selected with puromycin and confirmed by PCR for SV40 expression (Applied Stemcell). 293T fibroblast lines were maintained in DMEM supplemented with 10% FBS, penicillin/streptomycin, L-glutamine and sodium pyruvate. Transient transfections of wild-type or mutated MCM10-expressing plasmids were performed by transfecting 2×10^6^ cells with 6 μg of DNA using Lipofectamine 2000 according to manufacturers’ instructions (Invitrogen).

293T S-protein tag, Flag tag, Streptavidin binding peptide (SFB)-MCM10 stable cell lines were grown in DMEM (Gibco 11995) supplemented with 10% FBS (Sigma F4135), 1% penicillin/streptomycin (Gibco 15140) at 37°C and 5% CO2. Immortalized retinal pigmented epithelial cells (hTERT-RPE-1) were grown in DMEM:F12 supplemented with 10% FBS (Sigma F4135), 1 % penicillin/streptomycin (Gibco 15140) at 37°and 5% CO2. To generate a homozygous cell line for the R426C exon10 patient variant, a guide RNA targeting exon 10 of *MCM10* and a single stranded oligo containing the desired point mutation were transfected into HTERT-RPE-1 RPE-1. Cells were then subcloned and individual clones were screened for targeting. Biallelic knock-in of the mutation was confirmed with sequencing of TOPO TA clones. In a similar manner, heterozygous R582X (exon 13) patient MCM10 variant cell lines were generated using a guide RNA to the intronic region adjacent to the mutation location. A plasmid containing the point mutation and flanking regions of homology was used as a donor molecule to insert the patient *MCM10* variant. Cells were then subcloned and individual clones were screened for targeting. Biallelic knock-in of the mutation was confirmed with sequencing of TOPO TA clones.

### Plasmids

For generation of 805X, 771X, 705X and 607X truncation mutations in 293T cells *MCM10* coding sequence in the pDONR201 (Invitrogen) was used to generate truncation mutants using the QuikChange Lighting site-directed mutagenesis kit (Agilent). The full length and truncation sequences were transferred into a Gateway compatible destination vector that includes an N terminal triple-epitope tag (S protein tag, FLAG epitope tag and Streptavidin-binding peptide tag) using the LR clonase enzyme kit (ThermoFisher). Plasmids were transfected into 293T cells using FuGENE transfection reagent (Promega) and stable cell lines were selected using puromycin.

For generation of patient *MCM10* variants and expression in 293T cells full-length *MCM10* (NM_018518) was subcloned from pCMV6-X6 (Origene) into pLENTI-N-GFP. Site-directed mutagenesis was performed to generate missense (c.1276C>T, p.R426C) or truncation (c.1744C>T, p.R582X) mutations. 293T cells were transiently transfected using FUGENE6 and microscopy was performed 48 hours post-transfection.

### qPCR

RNA was extracted from WT or NK92 MCM10-KD cells using the RNeasy Mini kit (QIAGEN). First strand cDNA was generated using Superscript VILO MasterMix (ThermoFisher). qPCR for MCM10 was performed in quadruplicate on a Roche LightCycler with 25 ng of cDNA using TaqMan assay probe Hs00218560_m1 (ThermoFisher) or GAPDH as a housekeeping gene (Hs02786624_m1).

### Western blots

Immortalized fibroblasts from patient or healthy donor were lysed in RIPA buffer supplemented with Halt proteinase phosphatase (Pierce). For nuclear and cytoplasmic fractionation, cells were lysed in cytosolic buffer (10mM HEPES pH8, 10mM KCl, 0.5M EDTA pH 8, 0.2% NP40, 1X Halt proteinase phosphatase) and nuclear buffer (20mM HEPES pH8, 150mM NaCl, 0.5M EDTApH8, 1X Halt proteinase phosphatase). For co-immunoprecipitation of chromatin-associated proteins, 293T cells transiently transfected with p-Lenti turboGFP MCM10 plasmids were lysed in cytosolic buffer as described above, then the nuclear pellet was resuspended in co-IP buffer (10mM Tris-HCl pH7.5, 150mM NaCl, 0.5M EDTA, 0.5% NP40, 1X Halt proteinase phosphatase inhibitor) supplemented with 75U/ml DNase I and 2.5mM MgCl2 for 30’ at 4°C. Lysates were incubated with anti-turboGFP antibody (OTI2H8, Origene] or IgG isotype (NCG2B.01, Invitrogen) and Dynabeads Protein G (Invitrogen). Lysate for co-IP of CDC45 (D7G6, Cell Signaling Technologies) and MCM2 were incubated with turbo-GFP trap_MA antibody (Chromotek) or control binding antibody (bmab-20, Chromotek). For tight chromatin fractionation, cell pellets were subsequently washed in cytosolic buffer and nuclear buffer as described above and 50mM Tris-HCl pH8, 0.05% NP-40 buffers with increasing concentration of NaCl (0.15, 0.30M). Lysates were separated by gradient 4-12% gel and transferred to nitrocellulose membrane. Membranes were blocked using skim milk. Western blotting was performed using antibodies against the following: MCM10 (polyclonal, Proteintech), MCM2 (polyclonal, Abcam), POLA (polyclonal, Abcam), p53 (DO-1, Santa Cruz Biotechnology); lamin B1 (polyclonal, Abcam), α-tubulin (DM1A, Sigma); actin (polyclonal, Sigma). Secondary detection was performed using goat anti-Rabbit (Licor) or goat anti-mouse light chain specific (Licor) antibodies which were visualized by LiCor Odyssey and protein intensity was normalized to loading controls using ImageJ software. For 293T SFB-MCM10 stable cell lines, whole cell extracts were prepared as previously described (57) and analyzed with the following antibodies in 5% BLOT-QuickBlocker (G-Biosciences 786-011) in TBST: anti-FLAG antibody (Sigma F3165; 1:500 overnight at 4°C) and anti -α-tubulin (Covance MMS407R; 1:20,000 overnight at 4°C). Detection was with WesternBright Quantum detection kit (K-12042-D20).

For HTERT-RPE-1 mutant cells, whole cell extracts were prepared as previously described (57) and analyzed with the following antibodies in 5% BLOT-QuickBlocker (G-Biosciences 786-011) in TBST: anti-MCM10 (Novus Biologicals H00055388-D01P; 1:500 overnight at 4°C) which is a polyclonal antibody that was raised to full length MCM10 allowing detection of truncated protein, anti-MCM10 (Bethyl A300-131A; 1:500 overnight at 4°C) which was raised to an epitope in the C-terminal domain of MCM10, and tubulin (anti-α-tubulin, Covance MMS407R; 1:20,000 overnight at 4gC). Proteins were detected a with WesternBright Quantum detection kit (K-12042-D20).

### Doubling time assay

Cell doubling time was determined by plating 3 × 10^4^ cells per condition in a 6 well plate and counting 72 hours later. These conditions were established to not allow cells to become greater than 90% confluent during this assay to prevent contact inhibition from affecting doubling time analysis.

### Cell cycle analyses

For analysis of cell cycle, NK92 cells or primary patient fibroblasts were incubated with BrdU for 12 hours prior to fixation and staining with 7-AAD (BD Biosciences). Cells were analyzed via FACS on a BD Fortessa with 18-color configuration and resulting data were evaluated using Kaluza (Beckman-Coulter) or FlowJo (TreeStar) software. Where indicated, cells were pre-treated with aphidicolin to induce cell cycle arrest followed by washout. Quantification of cells found in different cell cycle stages was performed by gating analyses using the flow cytometric data.

### Cytotoxicity assays

NK92 or MCM10-KD cell lines were seeded at equal density prior to overnight culture to approx imate cell cycle synchronicity then were counted and incubated with 10^4^ K562 target cells that had been pre-incubated with 100 μCi 51Cr for 4 hours at 37°C 5% CO2. 1% IGEPAL (v/v) (Sigma-Aldrich) was used to lyse maximal release control wells and plates were centrifuged. Supernatant was transferred to a LUMA plate (Perkin Elmer) and dried overnight. Plates were read with a TopCount NXT and percent specific lysis was calculated as follows: (sample — average spontaneous release) / (average total release — average spontaneous release) × 100.

### Microscopy and image analysis

For preparation of microscopy samples, patient fibroblasts or transiently transfected cell lines were introduced into #1.0 Biotek imaging chambers and allowed to adhere overnight. Cells were fixed and permeabilized (Cytofix/Cytoperm, BD Biosciences) and incubated with polyclonal anti-MCM10 antibody (Abcam) followed by goat anti-rabbit secondary antibody conjugated to Alexa Fluor 488 (ThermoFisher). Cells were subsequently incubated with anti-*γ*H2AX antibody directly conjugated to Alexa Fluor 647 (2F3, Biolegend) and anti-fade mounting media containing DAPI was added to the wells. For visualization of 293T cells transiently expressing GFP-containing vectors expressing patient mutations, cells were grown in 8 well chamber slides with removable chambers for 24-48 hours after transfection. At this time, cells were fixed and permeabilized, then chambers were removed and coverslips were mounted with ProLong Gold antifade with DAPI (Thermo Fisher).

Images were acquired on a Leica SP8 confocal microscope equipped with a 100X 1.45 NA objective, with excitation by tunable white light laser. Detection was by HyD detectors and images were acquired with LASAF software. For some experiments, images were acquired on a Zeiss Axio-plan Observer Z1 with Yokigawa CSU-10 spinning disk and 63X 1.49 NA objective. Excitation was by 405 nm, 488 nm, 561 nm and 637 nm Coherent OBIS LX lasers controlled by a MultiLine Laserbank and controller (Cairn) and images were recorded with a Hamamatsu Orca R2 camera. Data were acquired by MetaMorph. After acquisition, all data were exported and analyses were performed in Fiji (58). For measurement of nuclear size, area of DAPI staining was ascertained. For measurement of nuclear MCM10 or GFP intensity, nuclei were segmented using DAPI and the intensity of MCM10 or GFP above negative control was quantified for each nucleus. For measurement of *γ*H2AX, the intensity and area of signal above negative control were measured. Data were graphed and statistical analyses performed in Prism 6.0 (GraphPad Software).

For 293T SFB-MCM10 stable cell lines, cells were plated on Millicell EZ slides (Millipore PEZGS0416) and allowed to recover for 24 hours. Cells were fixed with 3% paraformaldehyde in 1xPBS for 15 minutes and then permeabilized with 0.5% Triton-X-100 in 1xPBS for 5 minutes. Cells were stained with primary anti-FLAG antibody (Sigma F3165; 1:500) for 30 minutes at 37°C, followed by secondary antimouse Alexa Fluor 488 (ab150113), DAPI staining, and mounted using Vectashield (Vector Laboratories H-1000). Images were acquired using an Olympus FluoView FV1000 IX2 Inverted Confocal (University Imaging Centers at the University of Minnesota) and processed using FIJI and Adobe Photoshop.

For HTERT-RPE-1 mutant cell lines, cells were plated on #1.5 coverslips and allowed to recover for 24 hrs. Cells were then washed with PBS and treated with 20J UV and media replaced. For those cells that were untreated, they were washed with PBS and had fresh media immediately replaced. Cells were allowed to recover for 24 hours before being fixed in 3% paraformaldehyde in 1X PBS for 10 minutes and then permeabilized with 0.5% Triton-X-100 in 1xPBS for 5 minutes. Cells were blocked for 1 hour at room temperature in ABDIL (20mM Tris pH 7.5, 2% BSA, 0.2% fish gelatin, 130mM NaCl) followed by staining with primary anti-*γ*H2AX (Bethyl A300-081A) in ABDIL overnight at 4°C. They were then washed 3 times in PBST (1x PBS and 0.1% Tween) before staining with secondary anti-rabbit Alexa Fluor 488 (Thermo Fisher) for 1 hour at room temperature. Coverslips were washed 3 times in PBST, where the second wash contained DAPI, and mounted using Vectashield (Vector Laboratories H-1000). Slides were imaged on a Zeiss spinning disc confocal microscope using 100X 1.45 NA objective and 405nm and 488nm lasers. For image analysis Fiji was used to flatten images from both channels. After datasets were blinded, points with local maximal intensity were identified using Fiji and counted for each nucleus.

### In vitro NK cell differentiation

CD34^+^ HSC were isolated by FACS sorting from discarded apheresis collections from Texas Children’s Hospital. Initial lineage depletion was performed using RosetteSep enrichment (StemCell Technologies), followed by incubation with anti-CD34 PE antibody (4H11, eBiosciences), anti-CD56 (HCD56, Biolegend) and anti-CD3 (SK7, Biolegend). Cells were gated on the CD56^−^CD3^−^CD34^+^ population and collected. Following collection, CD34^+^ cells were nucleofected with the MCM10 CRISPR guide with a GFP tag described above or a GFP-expressing negative control, then maintained in IL-3, thrombopoietin, IL-6 and IL-7 for 3 days (44). Cells were sorted for GFP positivity and 2000 cells were co-cultured with EL08.1D2 stromal cells for 28 days in the presence of IL-3 (first week only), IL-6, IL-7, Flt3L, SCF and IL-15 (44). Following co-culture, lymphocytes were analyzed by FACS for cell surface receptors associated with NK cell development. Subsets were quantified using previously described phenotypic markers that mark progression from Stage 1 (CD34^+^ CD117^−^ CD94^−^CD16^−^) through Stage 2 (CD34^+^CD117^+^CD94^−^CD16^−^), then Stage 3 (CD34^−^CD117^+^CD94^−^CD16^−^), Stage 4 (CD34^−^ CD117^+/−^ CD94^+^ CD16^−^) and Stage 5 (CD34^−^CD117^−^CD94^+/−^CD16^+^) (45).

### In vitro iPSC generation

Dermal fibroblasts were expanded and reprogrammed into iPSCs using a non-integration method based on modified mRNA encoding Oct4, Klf4, Sox2 c-myc, Lin28, and control GFP (59, 60) using StemRNA repogramming (StemGent). Newly derived iP-SCs were passaged at least three times before banking. Cell lines were evaluated by karyotype analysis using G-banded karyotyping. They were further characterized for expression of pluripotency genes using qPCR (REX1, SOX2, NANOG and HTERT) and immunofluorescence (TRA-1-60 (Millipore; MAB4360), SSEA-3 (Millipore; MAB4303), SSEA-4 (Millipore; MAB4304), NANOG (Abcam; ab21624) and OCT4 (Abcam; ab19857). Lastly iPSC lines were differentiated in a spontaneous differentiation assay and expression of markers of three germ layers was assessed using qPCR for GATA4, AFP, CTnT, FLK1, PAX6, TUBB3, BRA and TUJ1. They were further characterized by karyotype analysis, expression of pluripotency genes and differentiation genes in a spontaneous differentiation assay. RNA was extracted using RNeasy Mini-Kit (Qiagen) and complementary DNA synthesized using KAPA cDNA Mix. iPSCs were maintaned on Matrigel in mTESR media at 37°C/5% CO2.

### In vivo HSC generation

To generate human HSC, 2 x 106 iPSCs were mixed with 106 OP9w3a that express WNT3a, and the cell mixture injected into female 4-8 week old NSG mice for teratoma formation (61): In order to create niche space for transplanted human hematopoietic stem cells, one day old NSG or Sirp1a pups received a sublethal dose of 100 cGy of X-ray irradiation in a biological irradiator. iP-SCs and Op9W3a cells were detached from culture flasks using 0.25% trypsin-EDTA when they reached 80% confluence. Cells were washed with PBS and resuspended in PBS + 2% FBS. Cells were counted and iPSCs were mixed with Op9W3a cells at a 2:1 ratio, and resuspended in PBS + 2% FBS so that the final volume was 100 *μ*L per mouse to be injected. NSG mice were injected with 2 million iPSCs and 1 million Op9W3a cells (3 million total cells in 100 *μ*L solution) subcutaneously into the dorsal neck region.

### Human CD34 HSC magnetic column purification and reconstitution of NSG mice

Eight to twelve weeks after injection of OP9w3a secreting and iPS cells, CD34^+^ HSC were isolated from teratomas of recipient NSG mice. A 1 mg/mL solution of Collagenase/ Dispase (Sigma) was prepared and injected into multiple points of teratomas to ensure penetration into the tissue. Teratomas were then minced into 1-mm pieces and poured into 15-mL conical tubes with approximately 10 mL of collagenase/dispase enzyme solution per teratoma. The tubes were incubated for 90 minutes at 37°C on a platform agitator. After incubation, the teratomas were filtered three times through 70 *μ*M nylon mesh and washed with PBS + 2% FBS. The teratoma cell suspension was then poured onto a Ficoll gradient and centrifuged for 20 minutes, 300g at room temperature. The cell layer was extracted and counted to prepare for magnetic column purification. A Miltenyi Biotec MACS CD34 MicroBead kit was used to positively select hematopoietic stem cells from teratoma single cell suspension after magnetic column purification. Cell suspensions were incubated for thirty minutes at 4°C with FcR blocking reagent and magnetic microbeads bound to anti-human CD34 antibodies. The solution was then applied to a positive selection column, which was held in place by a magnet. Buffer containing PBS, 0.5% bovine serum albumin, and 2 mM EDTA was used to flush columns. The column flowthrough was discarded, and the eluent was applied to a second column to increase purity. The final eluent was counted and a small aliquot was used to check % purity of HSCs via flow cytometry. Purity was measured as % of CD34^+^ cells out of total cells and was generally above 97%. The cells were then washed and resuspended in PBS + 2% FBS to achieve a final concentration of 50,000 HSCs per 20 μL. Previously irradiated one-day-old pups were injected intrahepatically with 50,000 HSCs, and were monitored routinely for survival and development of graft versus host disease.

### Isolation of immune cells from humanized mice for flow cytometric analysis of NK cells

Blood: Heart bleeds with a syringe were performed on mice immediately after death. Blood was collected in a FACS tube containing approximately 500 *μ*L PBS + heparin. Blood samples were diluted 1:1 with PBS and layered over a Ficoll gradient. The gradients were centrifuged for 20 minutes, 300g at room temperature. The cell layer was extracted and counted to prepare for flow cytometry staining. Spleen: Frosted slides were used to disaggregate the spleens into a solution of PBS + 2% FBS. Cells were filtered through 40 *μ*M nylon mesh and centrifuged for 5 minutes, 300g at room temperature. Red Blood Cells were lysed using Ammonium-Chloride-Potassium buffer for 2 minutes followed by washing of the cells with PBS + 2% FBS and centrifugation for 5 minutes, 300g at room temperature. The volume was brought to a convenient mark for counting, and the cells were counted to prepare for flow cytometry staining. Bone marrow: Right and left tibias and femurs were removed from mice during dissection. A syringe loaded with PBS + 2% FBS was used to flush out the bone marrow. This solution was filtered through 40 *μ*M nylon mesh and centrifuged for 5 minutes, 300g at room temperature. The volume was brought to a convenient mark for counting, and the cells were counted to prepare for flow cytometry staining.

### Analysis of NK cells from humanized mice by flow cytometry

Cells were isolated as described above and kept on ice for all immunostaining. Human NK cells were detected using the following antibodies: CD57 (Pacific Blue, clone HCD57, Becton Dickinson), CD56 (Brilliant Violet 605, clone HCD56, Biolegend), CD45 (PE-Cy5, HI30, Biolegend), CD3 (Brilliant Violet 711, clone SK7, Biolegend), CD62L (Brilliant Violet 655, DREG56, Biolegend, CD16 (PE Texas Red, clone 3G8, BD Biosciences), CD94 (APC, DX22, Biolegend), CD34 (PE, 4H11, eBiosciences), CD117 (PE Cy7, 104D2D1, Beckman Coulter). Incubation with antibody was performed on ice for 30 minutes prior to washing with PBS 2% FCS. Compensation controls were prepared using single stained OneComp beads (BD Biosciences). Fluorescence minus one controls were additionally prepared for CD45 and CD56 with splenocytes. Data were acquired on a BD Fortessa and analyzed in FlowJo (TreeStar). Approximately 10^6^ events were recorded from each organ from 4 mice per genotype.

## ACKNOWLEDGEMENTS

This work was supported in part by NIH R01AI120989 to JSO, NIH R01AI37275 to EMM, NIH GM074917 to AKB, NIH T32-CA009138 (R.M.B.), NIH National Cen-terforAdvancing Translational Sciences TL1R002493 and UL1TR002494 (M.M.S.) and the University of Minnesota Imaging Centers. JRL is supported in part by the National Institutes of Health NINDS (R35NS105078) and NHGRI/NHBLI (UM1 HG006542) to the Baylor Hopkins Center for Mendelian Genomics. The authors wish to acknowledge Mr. Blake Heath, Mr. Jansen Smith, Dr. Laura Angelo and Ms. Emily Haines for technical assistance with the development and experimentation on the humanized mouse model and Dr. Diane Yang for technical assistance with induced pluripotent stem cell generation and validation.

## AUTHOR CONTRIBUTIONS

EMM, JSO, AKB, SP and JRL designed experiments and interpreted data. EMM, JSO wrote the manuscript. MIC, RMB, MS, ST, AET, NCG, MM, and PPB performed experiments and analyzed data. The exome sequencing and interpretation of data were performed as part of the immunodeficiency cohort testing at BHCMG, and the proband in the current publication is designated individual number 68.1 in (53), where MCM10 was designated “potential novel gene 4”. IKC, ZCA, SNJ, A S-P, RAG, and DMM performed exome sequencing and exome data analyses. SJ, PPC, AGH and CS guided clinical evaluation of the patient and supervised clinical care. SP conceptualized, designed and performed experiments for reconstitution of NSG mice with patient iPSCs. MB and JC designed, performed and validated reprogramming of patient cells to iPSCs.

## CONFLICT OF INTEREST

Baylor College of Medicine (BCM) and Miraca Holdings Inc. have formed a joint venture with shared ownership and governance of the Baylor Genetics (BG), which performs clinical microarray analysis and clinical exome sequencing. JRL serves on the Scientific Advisory Board of the BG. JRL has stock ownership in 23andMe, is a paid consultant for Regeneron Pharmaceuticals and is a coinventor on multiple United States and European patents related to molecular diagnostics for inherited neuropathies, eye diseases, and bacterial genomic fingerprinting.

---

**Supplemental Figure 1.**
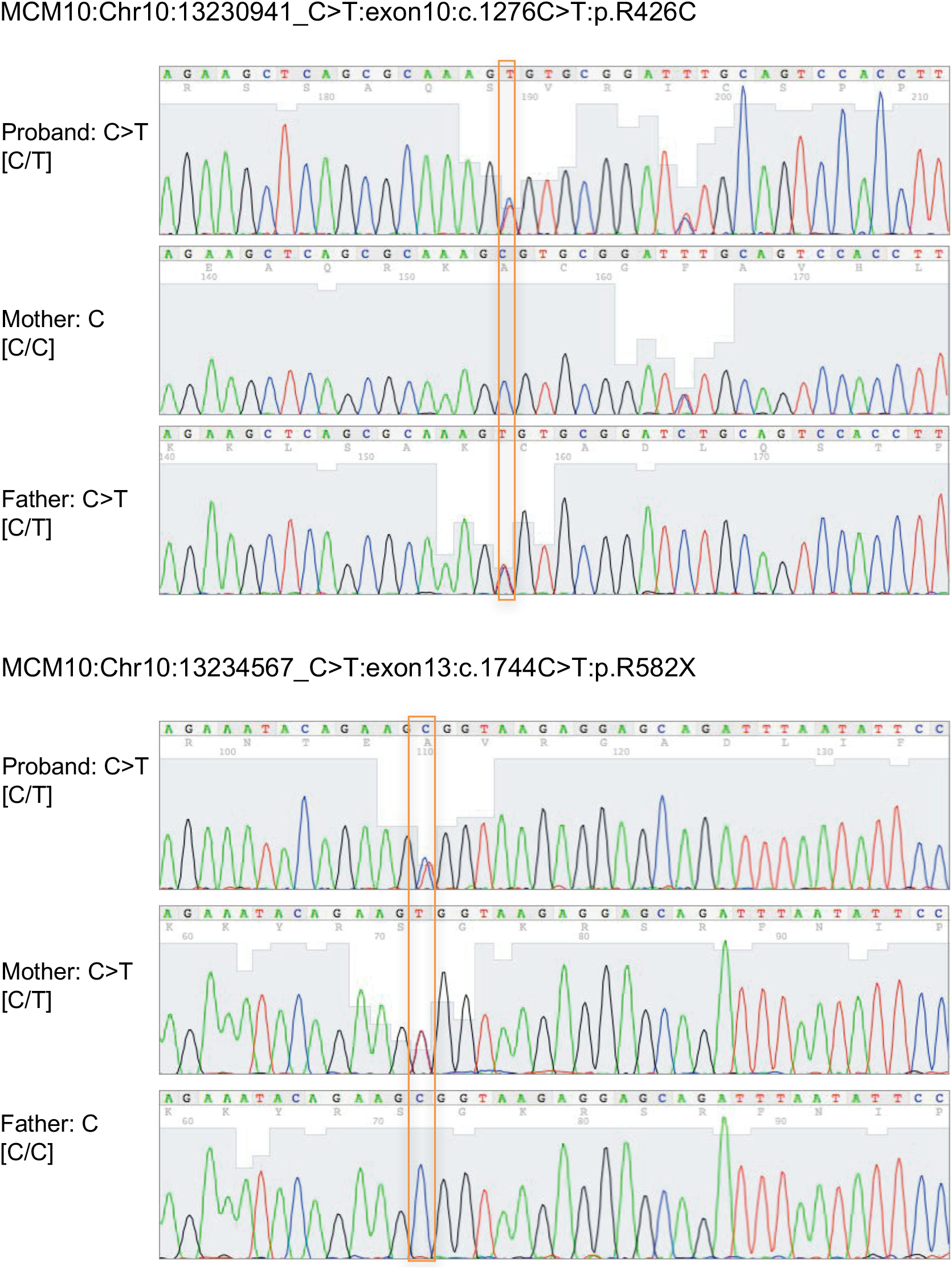
Sanger sequencing confirmation of c.1276C>T and c.1744C>T nucleotide variants in the patient and his parents (GRCh37/hg19).

**Supplemental Figure 2.**
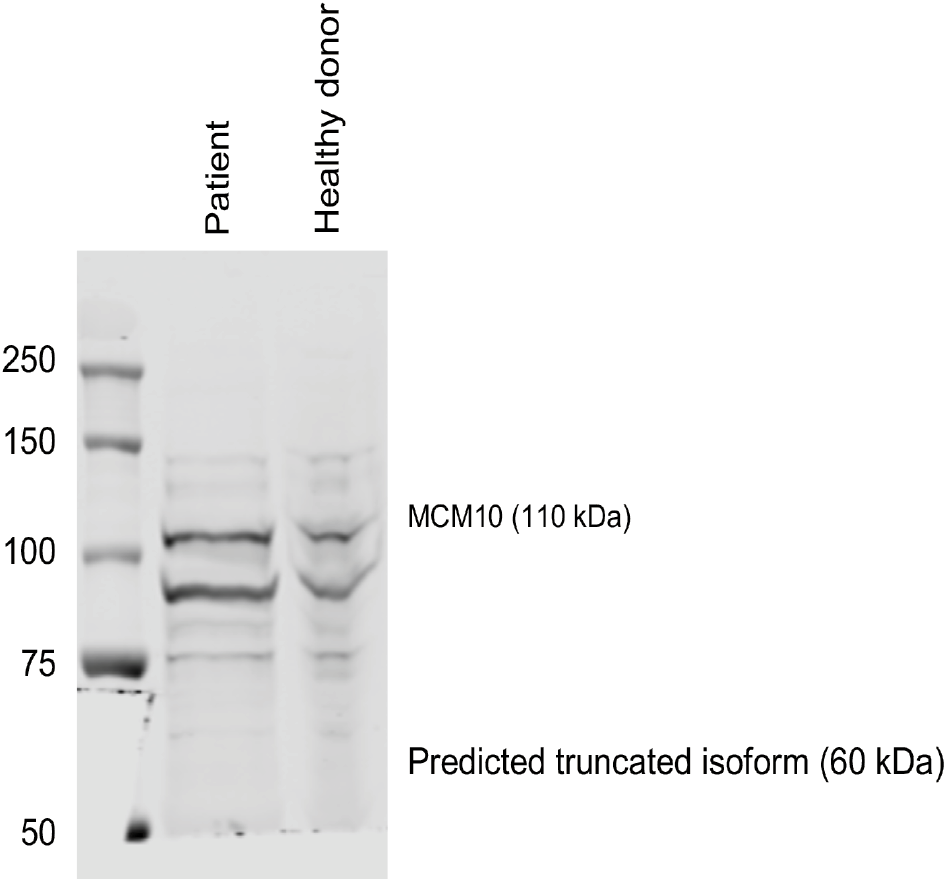
Western blot for MCM10 using full-length polyclonal immunogen. A) Primary fibroblasts from the proband and a healthy donor were lysed and probed for MCM10 protein (left) using polyclonal antibody raised against full-length MCM10. Predicted MW of truncated isoform=60 kDa.

**Supplemental Figure 3.**
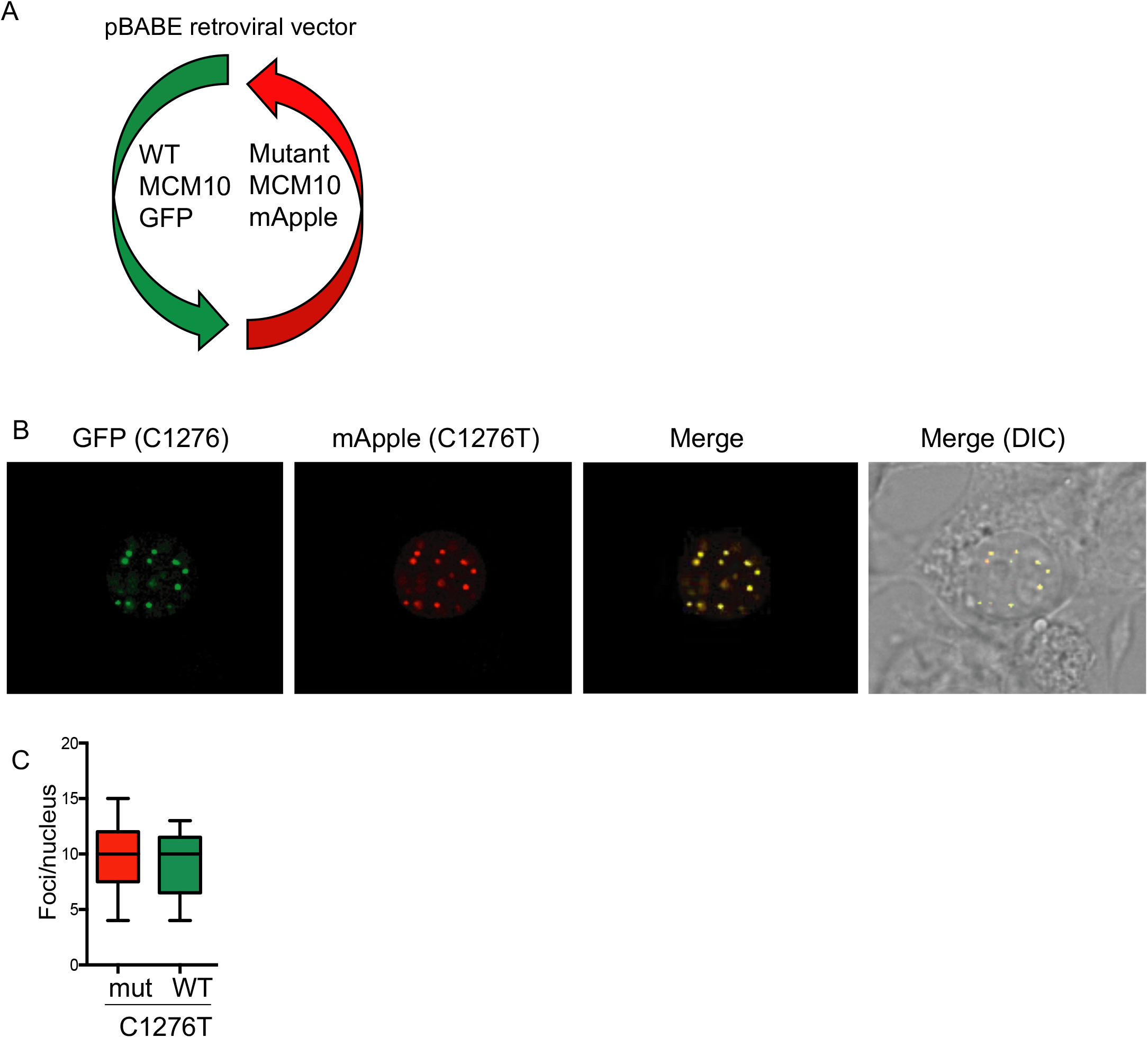
Bicistronic WT/R426C vector. A) Schematic of pBABE vector expressing full-length WT MCM10 with a GFP reporter, and MCM10 R426C with an mApple reporter. B) Transient transfection of the bicistronic vector in Phoenix cells followed by live cell imaging. C) Fluorescent foci per nucleus were enumerated using FIJI.

